# Functional Network Analysis of Fungal Pathogen *Colletotrichum sublineola* Effectors in *Sorghum* Anthracnose

**DOI:** 10.64898/2026.03.07.710159

**Authors:** Claudia Lerma-Ortiz, Janaka N. Edirisinghe, Purbasha Nandi, Clint Magill, Danyka Ramos-Melendez, Qun Liu, Christopher S. Henry

## Abstract

*Colletotrichum sublineola* (Cs) is a hemibiotrophic fungal pathogen that causes anthracnose in *Sorghum bicolor*, leading to significant yield losses. To enable infection, Cs secretes effectors - proteins, small RNAs, and metabolites - that damage the plant cell wall or enter the plant cell to suppress immune responses and manipulate host metabolism. Effectors can detoxify host antimicrobials, alter nutrient processing, and evade host immunity. Paradoxically, some effectors can also trigger pattern-triggered immunity (PTI), especially in biotrophic and necrotrophic fungi. More than half of fungal protein effectors lack conserved domains and functional network annotations. In this study, we identified prospective Cs effectors, separating those with non-conserved domains and classifying those with conserved domains by protein families. Comparative genomics is employed to predict effector functions and analyze their roles. Using their predicted locations and domains, we mapped the effectors into functional subsystems related to PTI. These include interactions in the apoplast, oxidative stress response, protein modification and degradation systems, and <u>C</u>ysteine-rich <u>F</u>ungus-specific <u>E</u>pidermal Growth Factor-like <u>M</u>odule (CFEM) domain proteins involved in immune regulation. Our functional network analysis advances the understanding of Cs pathogenicity and offers insights into effector infection mechanisms.

## Introduction

*Colletotrichum sublineola* (Cs), the causal agent of anthracnose in *Sorghum bicolor*, can reduce grain yield by up to 80% in susceptible cultivars (<u>Mekonen M et al., 2024</u>). Cs is a hemibiotrophic fungus (<u>Abreha KB et al., 2021</u>), which infects via biotrophic and necrotrophic stages. It secretes various infection agents called effectors—proteins, metabolites, and small RNAs—that translocate to the plant apoplast or cytoplasm, contributing to pathogenesis (<u>Abreha KB et al., 2021</u>; <u>Li G et al., 2024</u>; <u>Vela S et al., 2024</u>).

Cs protein effectors include “bona-fide” ones, small proteins with non-conserved domains (<u>Sonah H et al., 2016</u>; <u>Li G et al., 2024</u>; <u>Le Naour-Vernet M et al., 2025</u>) and proteins with conserved domains that serve enzymatic or binding functions (<u>De Wit PJGM et al., 2016</u>). Notably, some domain-conserved proteins retain structural features, enabling them to suppress host immunity despite lacking enzymatic activity (<u>Fiorin Gl et al., 2018</u>). Effectors that have non-conserved domains are small, rapidly evolving proteins which enhance fungal pathogenicity. In contrast, certain effector families with conserved domains exhibit limited variation and are widely distributed among pathogenic fungi. These conserved domain features enable researchers to: (1) predict effector functions through comparative genomics to guide experimental validation, and (2) identify targets for engineering plants with improved resilience. For instance, plants have been engineered to counteract fungal polygalacturonase and toxins such as eutypine, fumonisin B1, deoxynivalenol, and cercosporin (<u>Punja ZK, 2006</u>). Apoplastic effectors are synthesized in the endoplasmic reticulum (ER) and secreted via the ER-Golgi pathway. In contrast, cytoplasmic effectors utilize alternative secretory routes, influenced by their codon usage bias (<u>Li G et al., 2024</u>). Apart from secretion signal peptides, cytoplasmic effectors often bear additional motifs—such as transit peptides or nuclear localization signals—that direct them to host organelles like chloroplasts, mitochondria, or the nucleus (<u>Sperschneider J et al., 2017</u>), though exceptions exist (<u>Nielsen H et al., 2019</u>). These structural features inform predictive tools, including <u>SignalP 6.0</u> for secretion signals (<u>Nielsen H et al., 2019</u>), <u>EffectorP 3.0</u> for effector identification, and <u>Localizer</u>, for host organelle targeting (<u>Sperschneider J et al., 2017</u>; <u>Sperschneider J and Dodds PN, 2022</u>).

A complex cell wall comprising a primary wall, middle lamella, and a three-layered secondary wall cover the epidermal cells of *Sorghum bicolor* leaves. A cuticle overlays the wall and there is a deposit of cutin among the cellulose microfibrils of the outer wall layer. All these structures protect *S. bicolor* leaves against invasion. So initially, Cs forms an appressorium that adheres to the plant cuticle and secretes apoplastic enzymes and effectors to breach its cell wall (<u>Wharton PS et al., 2001</u>). It simultaneously forms a turgid penetration peg. During biotrophy, a primary hypha (haustorium) develops without breaching the host membrane or inducing cell death, instead suppressing pattern-triggered immunity (PTI) via effector secretion (<u>Abreha KB et al., 2021</u>). In the necrotrophic phase, secondary hyphae arise, invade host tissues, and induce cell death. The infection ultimately spreads to the rachis, panicle branches, and seeds, significantly reducing yield (<u>Vela S et al., 2024</u>). Fungal effectors participate in all these invasion stages, so it is imperative to determine their function and interactions to understand and combat fungal pathogenesis. In this study, we do a functional prediction of query Cs effector proteins informed by a comprehensive analysis of:

A. their amino acid sequences, which reveals several key features: (1) the presence of N-terminal signal peptides, indicative of secretion potential (<u>Nielsen H et al., 2019</u>); (2) an elevated proportion of positively charged residues and cysteine enrichment, which may influence subcellular localization within the host (<u>Sperschneider J and Dodds PN, 2022</u>); and (3) the identification of conserved domains through CDART-based homology searches, enabling inference of primary biochemical roles based on domain architecture (https://www.ncbi.nlm.nih.gov/Structure/cdd/cdd_help.shtml
B. experimental data available in the literature for homologous proteins. To refine functional predictions based on amino acid sequences, we validated and adjusted our predictions by employing the proposed function as a query term in AI type-assisted searches for experimentally characterized effectors, followed by comparative sequence homology assessments between these reference effectors and the query proteins. Collectively, these attributes allow for the prediction of an effector’s principal biochemical function—whether enzymatic or structural. Nevertheless, cellular components rarely operate in isolation; rather, they function within intricate physiological networks. This paradigm applies to fungal effectors, which orchestrate and modulate pathogenicity in coordination with fungal invasion strategies and host immune responses, including pattern-triggered immunity (PTI). Hence, we also consider these interactions in our subsystems design. We conclude by evaluating the strengths and limitations of comparative genomics in the study of fungal pathogenicity, particularly in the case of Cs.

## Methods

### Colletotrichum sublineola’s genome assembly

*C. sublineola* effector mapping across close genomes including the FSP237, Sequencing data, assemblies, close genomes, orthologous protein families are available at KBase Narrative https://narrative.kbase.us/narrative/172651

### Prediction of Cs effectors

The proteome of *C. sublineola* (UP000027238) containing 12,697 unique proteins was downloaded from Uniprot (https://www.uniprot.org). We identified candidate secreted proteins by predicting signal peptides using SignalP 6.0. We employed SignalP 6.0 which predicts the presence of signal peptides as well as their cleavage sites in proteins (https://www.nature.com/articles/s41587-021-01156-3). SignalP 6.0 identified 1428 protein sequences with signal peptides. To further predict the localization of these proteins, We passed the sample protein sequences to Wolf-PSORT (https://academic.oup.com/nar/article/35/suppl_2/W585/2920788?login=false) to predict their subcellular localizations. Wolf-PSORT analysis resulted in 1370 sequences with at least one homolog having confirmed extracellular localization. Finally, we applied EffectorP 3.0 (https://apsjournals.apsnet.org/doi/10.1094/MPMI-08-21-0201-R) to predict both apoplastic and cytoplasmic effectors. In total EffectorP 3.0 (https://effectorp.csiro.au/) predicted 410 effectors. Among them, 266 are apoplastic effectors, 99 are cytoplasmic effectors, and 45 are effectors with dual localizations to both apoplast and cytoplasm.

### Classification and comparative genomics

We used the NCBI Protein Domains and Macromolecular Structures platform (https://www.ncbi.nlm.nih.gov/Structure/cdd/wrpsb.cgi) to analyze the conserved domain content of the effector proteins. Based on these results, we divided effectors into proteins with conserved or non-conserved domains; the latter will be addressed in a separate study. Here, we classify conserved-domain effectors based on their common structures, enzymatic activities and location into functional groups in the context of fungal pathogenicity. We delineated four host-pathogen interaction subsystems: (1) apoplastic immune interplay, (2) oxidative stress response, (3) protein modification, folding, and degradation, and (4) CFEM-mediated immune regulation. We applied comparative genomics to predict effector functions and examine their roles within these frameworks. To do this, based on the conserved-domain composition of the Cs proteins, we generated a query description that captured terminology associated with their protein-family functions and combined it with targeted clue phrases to construct a functional descriptor for our AI-based search. This search typically yielded two or three principal effector candidates whose functions have been described in the scientific literature. We then compared the Cs proteins (subjects) with the proteins from other organisms (sources) identified in the search, examining sequence homology (including cross-BLAST results), structural features, and conserved-domain content. Finally, we analyzed the function of the source organism’s effector within its pathogenicity mechanisms.

## Results

This publication tables can be viewed and downloaded from this figshare webpage: https://doi.org/10.6084/m9.figshare.31347355

Among the 45 dual-localized effectors, 35 lack conserved domains, while the others include binding proteins (4), CAZymes (2), and one each of a virulence factor, aminotransferase, RNase, and DUF6413 (unknown function) (**Table 1_Dual_Location_Tab**). We identified 99 cytoplasmic effectors. Among them 34 have non-conserved and 65 have conserved domains **(Figure 1 and Table1_Cytoplasmic _Tab).** Additionally, we identified 266 apoplastic effectors, including 133 lacking conserved domains **(Figure 2 and Table1_Apoplastic_Tab).**

**Figure 1.**
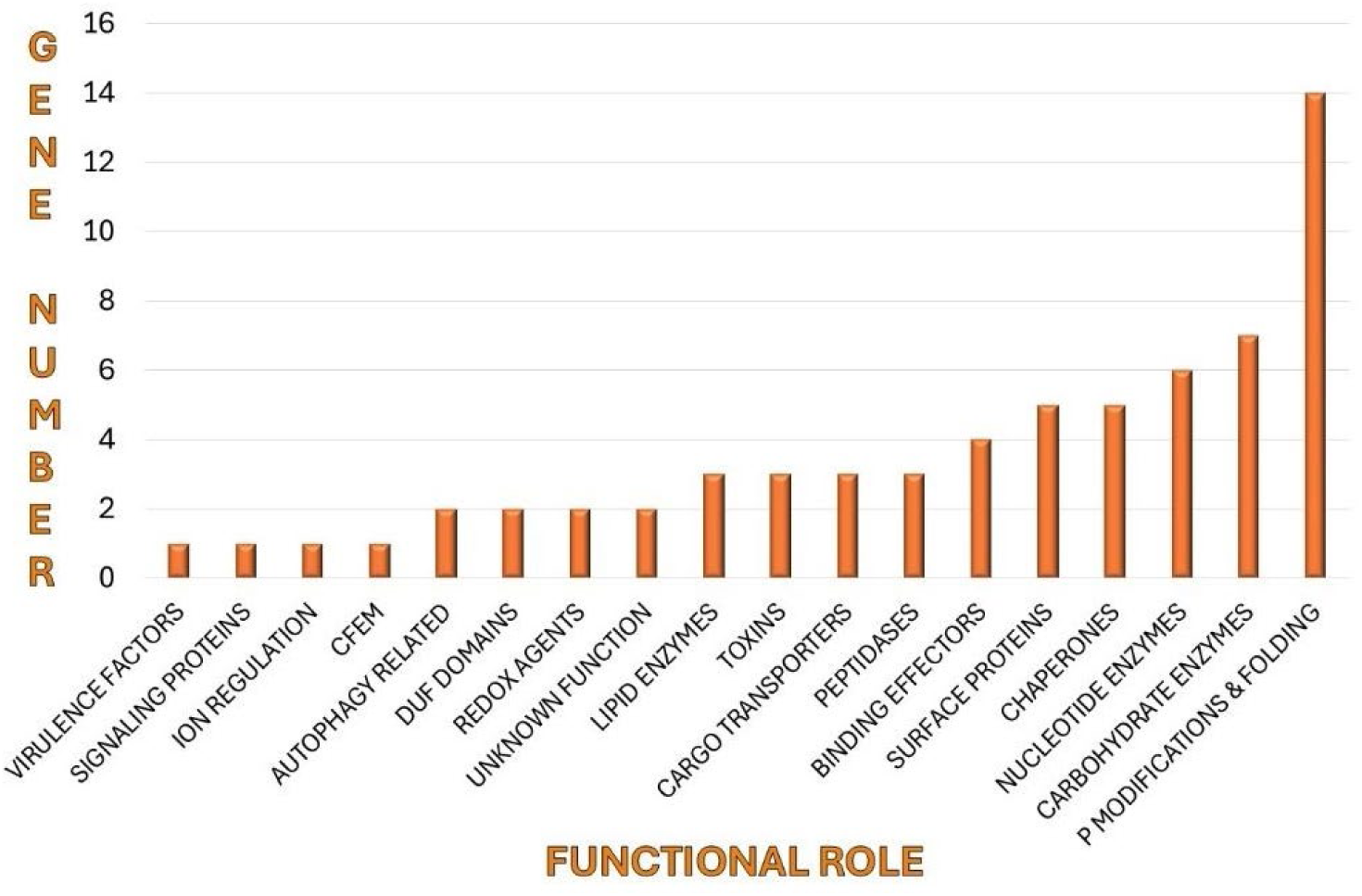
Functional distribution of *Colletotrichum sublineola* cytoplasmic effectors with conserved domains. Most of these domains were linked to protein modifications as discussed in this study. Effectors were identified using SignalP 6.0 and EffectorP 3.0, then classified by protein families via the NCBI Conserved Domains Database **(Table 1_Cytoplasmic_Tab).**

**Figure 2.**
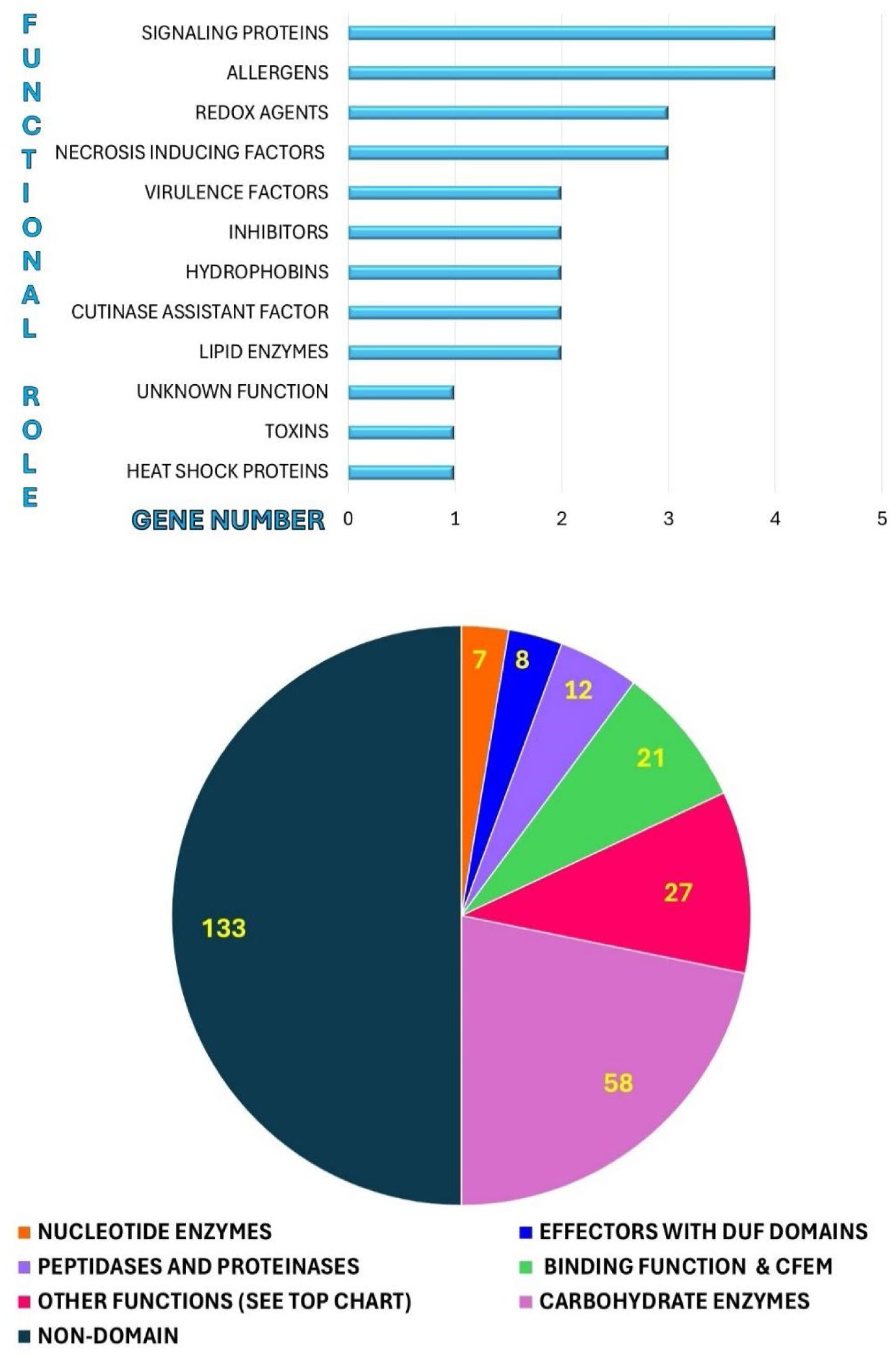
Functional distribution of *Colletotrichum sublineola* apoplastic effectors with conserved domains. The bottom pie chart displays the classification of the *C. sublineola* 266 apoplastic effectors, including 133 proteins with non-conserved domains, which have limited homology across taxa. Among apoplastic effectors with characterized domains, most (58) are CAZymes. Other effectors with enzymatic activity are the peptidases and proteinases (12) and the ones acting on nucleotides (7). Only eight proteins contain DUF domains of unknown function. The 27 effectors with various other functions are described on the top bar chart (see **Table1_Apoplastic_Tab** for details).

### Effectors Functional Networks

#### Network I) Apoplast Immune Mechanisms Interplay

Here we describe Cs apoplastic effectors, their diverse functions and how they collectively promote Cs pathogenicity.

##### Carbohydrate-Active Enzymes (Cazymes)

Fungi produce distinct CAZymes across their growth stages, and these enzymes networks have been extensively studied (<u>Samal A et al., 2017</u>). Here, we focus on Cs CAZymes with dual roles as effectors. The CAZyme effectors targeted to the apoplast can be classified into four groups: (1) glycosyl hydrolases, (2) monooxygenases, (3) pectinases, (4) other CAZymes (**Table 1_Apoplastic_effectors, and _Cytoplasmic_effectors_Tabs).** Their catalytic activities are detailed in **Table 2 (Glycosyl hydrolases, and Other_Apoplastic_Enzymes_Tabs**) and **Figure 3**.

**Figure 3.**
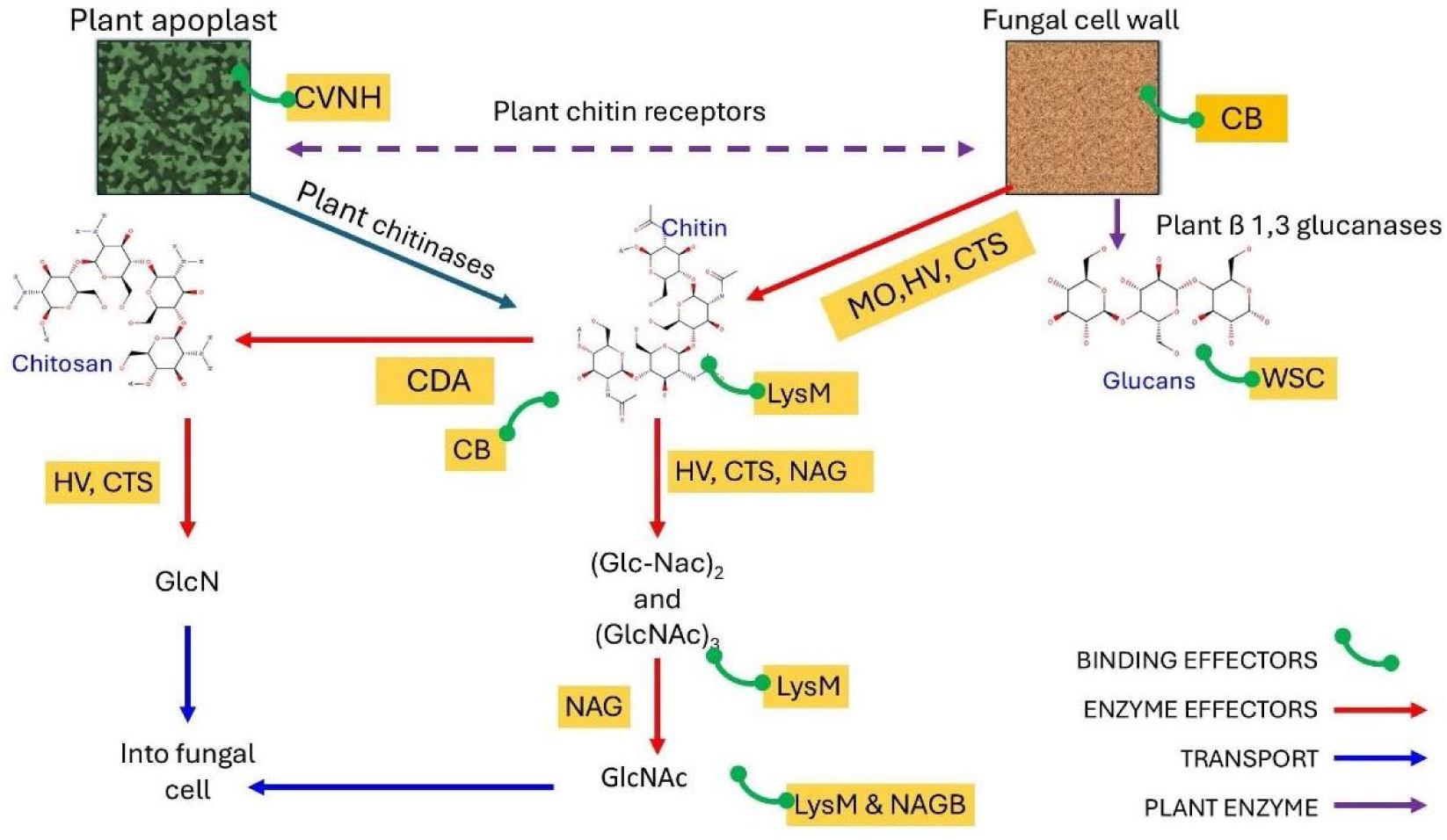
Functional interplay of effectors in the apoplast. A total of 21 Cs genes encode apoplastic effectors with binding functions, primarily targeting chitin and glucan degradation products. These binding proteins act synergistically with effector enzymes to prevent elicitation of Sorghum’s pattern-triggered immunity (PTI). A full list of apoplastic enzyme effectors, including CAZymes is shown in **Table1** and the Cs effectors with binding functions are classified in **Table 2_Binding_functions_Tab**. Abbreviations: MO, monooxygenase; HV, hevamine; CTS, chitinase; CB, chitin-binding domain; NAGB, N-acetylglucosamine-binding; CDA, chitin deacetylase; NAG, N-acetylglucosaminidase; CVNH, CyanoVirin-N; LysM, lysin motif; WSC, cell wall integrity and stress response component.

###### Glycosyl hydrolases

Among these, glycosyl hydrolases (GHs) act both in carbohydrate metabolism and as virulence factors, a dual role discussed further below. CAZymes contribute to cell wall degradation, chitosan remodeling, and immune evasion by interfering with host receptor detection.

Identified GH families include: GH7 (endo-β-1,4-glucanases; EC 3.2.1.4), GH10 and GH11 (endo-β-1,4-xylanases; EC 3.2.1.8), and GH12 (xyloglucan-specific endo-β-1,4-glucanases; EC 3.2.1.151; GH16 includes enzymes with diverse functions, such as endo-1,3(4)-β-glucanases (EC 3.2.1.6), BglS (COG2273) endo-β-1,3-galactanases (EC 3.2.1.181), and members of the cd02183 subfamily, which exhibit glycosylphosphatidylinositol-glucanosyltransferase activity.

Additional identified families include GH28 (endo-polygalacturonases; EC 3.2.1.15), GH43 (arabinan endo-1,5-α-L-arabinanases; EC 3.2.1.99), GH45, cellulases (EC 3.2.1.4), and GH75, chitosanases (EC 3.2.1.132). The precise activities of A0A066X6U4 and A0A066XI06—both GH16 members—remain unclear, though they likely belong to a subfamily capable of hydrolyzing glycosidic bonds between carbohydrates and between carbohydrate and non-carbohydrate moieties. We will cover the glycosyl hydrolase effectors with more detail later in this publication.

###### Monooxygenases

Cs encodes two types of monooxygenase effectors. First, thirteen LPMO_AA9 family members exhibit lytic cellulose monooxygenase activity—either C1-hydroxylating (EC 1.14.99.54) or C4-dehydrogenating (EC 1.14.99.56). Five of these proteins also have a CBM_1 cellulose-binding domain. Second, one AA13_LPMO effector contains dual domains: a lytic starch monooxygenase (EC 1.14.99.55) and a CBM20 glucoamylase (EC 3.2.1.3) **(Table 2_Other_Apoplastic_Enzymes_Tab)**. LPMO_AA9 enzymes act together with endoglucanases and β-glucosidases in degrading recalcitrant polymers (<u>Pierce BC et al., 2017</u>), while the AA13_LPMO effector targets starch, generating malto-oligosaccharides convertible to maltose (<u>Lo Leggio L et al., 2015</u>).

###### Pectinases

Cs has several glycosyl hydrolase family 28 effectors **(Table 2_Other_Apoplastic_Enzymes_Tab)** with potential pectin hydrolase activity, similar to that observed in *Aspergillus tubingensis* polygalacturonase (EC 3.2.1.15) (<u>Tai E-S et al., 2013</u>). Additionally, Cs encodes five pectate lyase (EC 4.2.2.2) effectors—three from pfam03211 and two from the PelB/COG3866 families. Effector A0A066XA72, a member of polysaccharide lyase family 1, likely exhibits pectate and/or pectin lyase activity (EC 4.2.2.10), while A0A066XBV1 is predicted to function as a pectin methylesterase (EC 3.1.1.11). Cs lacks homologs for pectate lyase 22 (EC 4.2.2.6), found in other fungi, and for bacterial-type pectate disaccharide-lyases (EC 4.2.2.9) (BRENDA database) **(Table 2 same Tab).**

###### Additional CAZymes and Other Apoplastic Enzymes

Some CAZymes exhibit bifunctionality by combining distinct glycosyl hydrolase domains. For instance, A0A066WZH1 from GH family 131 possesses both exo-β-1,3/1,6-glucanase (EC 3.2.1.58) and endo-β-1,4-glucanase (EC 3.2.1.6) activities, similar to *Coprinopsis cinerea* CcGH131B (<u>Shiojima Y et al., 2025</u>). Beyond glycosyl hydrolases, Cs has other CAZymes such as non-reducing end α-L-arabinofuranosidase (EC 3.2.1.55), glucan 1, 4-α-glucosidase (EC 3.2.1.3), and rhamnogalacturonan endolyase (EC 4.2.2.23) **(Table 2).** Cs also produces effectors like chitin deacetylase (EC 3.5.1.41) to process chitin degradation products, which arise from both plant defense and fungal cell wall remodeling during morphogenesis. Additionally, effectors such as cutinase (EC 3.1.1.74), which degrades wax type cutin, contribute to cuticle degradation.

##### Apoplastic Effectors with Binding Functionality

###### LysM-Containing Effectors

Plant cell walls contain peptidoglycan structures, which pathogenic fungi degrade using specialized enzymes to aid penetration. Many of these enzymes comprise separate catalytic and substrate-binding domains. One such domain, LysM (∼40 residues; NCBI domain database), features a βααβ secondary structure and binds peptidoglycans. Although of bacterial origin, the genes encoding for this structure likely entered eukaryotic genomes via horizontal gene transfer (<u>Bateman A et al., 2000</u>). LysM effectors are known to bind chitin, shielding it from host immune recognition (<u>Sánchez-Vallet A et al., 2020</u>; <u>Zeng T et al., 2020</u>). Cs encodes three LysM-containing effectors: A0A066XAG3, A0A066XAZ6, and A0A066X6X5. The first exhibits dual localization; the latter two are restricted to the apoplast. While A0A066XAG3 and A0A066XAZ6 have standalone LysM motifs, A0A066X6X5 integrates LysM within a 1,4-β-N-acetylmuramoylhydrolase enzyme **(Table 2_Binding_functions_Tab).**

###### CVNH Effectors

Some filamentous ascomycetes and ferns produce proteins with a <u>C</u>yano<u>V</u>irin-<u>N H</u>omology (CVNH) domain-homologous to cyanobacterial CVN lectins that bind mannoses with high affinity-. In fungi, CVNH domains occur in both secreted and non-secreted proteins, often within multidomain structures (<u>Pecurdani R et al., 2005</u>). Unlike cyanobacterial CVNs, fungal CVNHs lack <u>conserved</u> sugar-binding sites and cysteines, making their functions harder to predict. However, they retain carbohydrate-binding motifs with variable locations and affinities. For instance, *Neurospora crassa* and *Tuber borchii* CVNHs each have a single binding site in different domains, while *Ceratopteris richardii* has two (<u>Gronenborn AM, 2009</u>). *Colletotrichum sublineola* (Cs) encodes six standalone CVNH-domain effectors: A0A066XKV0 and A0A066X3D8 (apoplastic); A0A066XM58 and A0A066X047 (cytoplasmic); A0A066X0R6 and A0A066X047 (dual localization).

###### WSC Effectors

The “<u>C</u>ell <u>W</u>all Integrity and <u>S</u>tress Response <u>C</u>omponent” (WSC) domain, approximately 90 amino acids in length with eight conserved cysteines (<u>Verna J et al., 1997</u>), binds xylans and fungal chitin/β-1,3-glucans (<u>Wawra S et al., 2019</u>). In some fungal enzymes, including β-1,3-exoglucanases and alcohol oxidases, WSC domains serve as anchoring modules to fungal or plant cell walls (<u>Oide S et al., 2019</u>). Cs encodes six apoplastic effectors with one to three WSC domains (A0A066X7P7, A0A066XAP8, A0A066XC12, A0A066XLE7, A0A066XHU5, A0A066XIJ7). As these effectors lack additional domains, they are likely involved in chitin and glucan binding to mask fungal structures from plant immune receptors.

###### Chitin and Chitin-Derivative Binding Proteins

Additional chitin-binding architectures include (1) the Hevein (ChtBD1; cl16916) domain, which recognizes and binds chitin subunits, eg N-acetylglucosamine (https://www.ebi.ac.uk/interpro/entry/cdd/CD11618/); and (2) pfam00187 chitin-binding domains, typically fused to catalytic domains of enzymes such as plant chitinases (https://www.ebi.ac.uk/interpro/entry/InterPro/IPR001002/). These domains are cysteine-rich (<u>Wright HT et al., 1991</u>). The Cs effector A0A066Y1P5 contains a tandem of two chitin-binding domains (pfam00187) and a Hevein domain. Other types of proteins that also have conserved cysteines and bind N-acetylglucosamine oligosaccharides and chitin are the cerato-platanins (<u>Gaderer R et al., 2014</u>). Cs effectors A0A066X9M4 and A0A066XDH8 belong to this family.

###### Cellulose-binding proteins

The fungal cellulose binding (CBM_1) domain (pfam00734) contains four conserved cysteines forming disulfide bonds and adopts a three-stranded antiparallel β-sheet structure (https://www.ebi.ac.uk/interpro/entry/profile/PS51164/). It is typically fused to CAZymes with catalytic domains, such as endoglucanases (EC 3.2.1.4), exoglucanases (EC 3.2.1.8), and cellobiohydrolases (EC 3.2.1.91). In Cs, CBM_1 is fused to an acetyl xylan esterase AxeA (EC 3.1.1.72, (A0A066WYK6) and to several monooxygenases (A0A066XRP6, A0A066XNJ6, A0A066XNR3). Notably, protein A0A066X146 in Cs functions as a standalone CBM_1 **(Table 2_Binding_functions_Tab)**.

#### Functional Interplay of Effectors in the Apoplast

Fungal effector secretion varies by infection stage, aligning with the pathogen’s physiological goals. These effectors act synergistically with each other and host systems. Expression profiling and experimental data can help identify effectors active during Cs pathogenesis. Cs employs four combinatorial effector strategies: (1) chemical and physical disruption of the plant cell wall; (2) amplification of this attack; (3) masking of fungal presence; (4) the vanishing game (evasion).

##### Chemical and physical disruption

*The appressorium penetration:* to breach the cuticle, the appressorium secretes adhesive proteins, including hydrophobins (<u>Wang Z et al., 2010</u>). Cs produces two hydrophobin type 2 effectors (pfam06766)—A0A066WYW8 and A0A066XKB8—known as cryparins, which are lectin-like proteins that adhere to the fungal cell wall and are secreted extracellularly (<u>McCabe PM and Van Alfen NK, 1999</u>). Appressoria develop high turgor pressure and release CAZymes to degrade plant barriers. This concerted physical and chemical attack facilitates the peg penetration through the plant cell wall.

##### Amplification of the attack

*Degradation boosters:* as described above, the combined activity of lytic cellulose monooxygenases, endoglucanases, and β-glucosidases markedly enhances plant cell wall degradation (<u>Pierce BC et al., 2017</u>) **(Figure 3).** Cs also produces two effectors, A0A066XF37 and A0A066XHT4, from the hydrophobic surface binding protein A family (pfam12296). These proteins are believed to adsorb to hydrophobic surfaces, such as the leaf cuticles, and recruit enzymes like cutinases, thereby facilitating the plant surface degradation (<u>Ohtaki S et al., 2006)</u>.

##### Masking Strategy

Binding-effectors function in combination with fungal enzymes, or other types of effectors or even plant enzymes, to prevent degradation products from activating plant pathogen recognition receptors, thereby suppressing pattern-triggered immunity (PTI) **(Figure 3).**

This operation consists of targeting the following metabolites:

Chitin and its Derivatives Masking: Fungal growth and invasion involve dynamic remodeling of the fungal cell wall, including chitin synthesis and degradation (<u>Malinovsky FG et al., 2014</u>). Cs secretes chitinases and corresponding degradation products. When the host plant (e.g., *Sorghum*) detects fungal cell walls on its surface, it also releases chitinases that liberate chitin oligomers into the apoplast. In response, Cs deploys multiple chitin-binding effectors—cerato-platanins, WSC, LysM, and heveins—that bind chitin fragments down to N-acetylglucosamine. Once sequestered by these effectors, the oligomers cannot engage plant immune receptors. Alternatively, fungi produce chitin deacetylases that convert it into chitosan. This polymer and its degradation products do not elicit PTI.

Masking Polysaccharide Degradation Products: Cs secretes WSC-type glucan-binding effectors that shield polysaccharides from host glucanases and recognition by plant receptors. Degradation of the *Sorghum* cell wall releases mannose and xylans (<u>Scavuzzo-Duggan T et al., 2021</u>), which are masked by CVNH (mannose) and WSC (xylans) effectors, respectively, preventing recognition by host pattern recognition receptors (PRRs).

##### The Vanishing Game

Chitosan can be further hydrolyzed into glucosamine by chitosanase-active effectors. Both glucose and N-acetylglucosamine are then imported into the fungal cell, effectively removing them from the apoplast.

#### Network II) Effector-Mediated Modulation of the Plant Oxidative Stress Response

Hydroxyl radicals (^⋅^OH) and superoxide (^⋅^O₂⁻) are highly reactive oxygen species (ROS) (<u>De Gara et al., 2010</u>). While hydrogen peroxide (H₂O₂) is less reactive, it generates these radicals upon interacting with metal ions (<u>Demidchik V, 2015</u>) and is thus also considered a ROS. Plants produce and accumulate ROS as part of defense and signaling responses to biotic and abiotic stresses (<u>Demidchik V, 2015</u>). To counteract ROS-mediated damage, fungal pathogens have evolved antioxidant enzymes, specialized metabolites, and effectors that neutralize ROS and interfere with ROS-dependent signaling (<u>Singh Y et al., 2021</u>). Effector type and timing depend on the fungal lifestyle—necrotrophic, biotrophic, or hemibiotrophic—and can either stimulate or suppress ROS production. In addition to signal peptides, effectors often possess targeting motifs that direct them to specific plant organelles, so they can be directed where they are most effective. Structural analysis of effectors, alongside knowledge of plant ROS-generating mechanisms, may help predict their roles in modulating oxidative stress. Key sources of ROS in plants include mitochondria, ER, chloroplasts, and peroxisomes. Under stress, additional ROS production arises from plasma membrane–localized NADPH oxidases (<u>Singh Y et al., 2021</u>), cell wall peroxidases, quinone reductases, and oxalate and amine oxidases (<u>Jiménez-Quesada et al., 2016</u>). The NADP-malic enzyme provides electrons to the NADPH oxidase as needed. The latter are located on the plasma membrane and transport electrons from a cytosolic electron donor to extracellular oxygen, generating superoxide (^⋅^O2-) which is rapidly converted to H_2_O_2_ spontaneously or by superoxide dismutase. Chloroplasts also produce ^⋅^OH radicals via both photosystems, contributing to signaling but posing a risk of oxidative organellar damage if not controlled (<u>Jwa NS and Hwang BK, 2017</u>; <u>Jiménez-Quesada et al., 2016</u>). Effector strategies for ROS suppression then include: (1) direct ROS scavenging; (2) inhibition of ROS producing enzymes; (3) modulation of host regulation of ROS-generating pathways.

The last mechanism is the most sophisticated one and hence the effectors that operate this way are difficult to discover.

##### Effectors Modulating ROS

**Table 3** shows some of the enzymes involved in hydrogen peroxide (H₂O₂) metabolism listed in the BRENDA database, including both producers and scavengers, some of which have been co-opted by fungi as effectors. These effectors are secreted into the plant apoplast or cytoplasm at various infection stages, depending on the pathogen’s strategy. **Table 1** includes peroxidase, cupredoxin (apoplastic) and glutaredoxin (cytoplasmic), as some of the redox effectors secreted by Cs, and **Table 4 (Oxidative_stress_Tab)** shows some homologs of these apoplastic effectors in other pathogens.

A0A066X6A6 is an apoplastic effector plant type peroxidase (EC1.11.1.7). Peroxidases use a wide range of aromatic compounds as electron donors, including melanin to catabolize H_2_O_2_, but when needed, they can also produce H_2_O_2_ by an oxidative cycle using reductants found in the apoplast (<u>Černý M et al., 2018</u>) **(Figure 4, top panel)**. A0A066X6A6 presents some similarity to BkLiP1, a peroxidase that acts mainly as virulence factor in *Botryosphaeria kuwatsukai* (E= 1.9 e-43 with 45.60% % identity), inducing ROS bursts and plant cell death (<u>Xiao F et al., 2022</u>).

**Figure 4.**
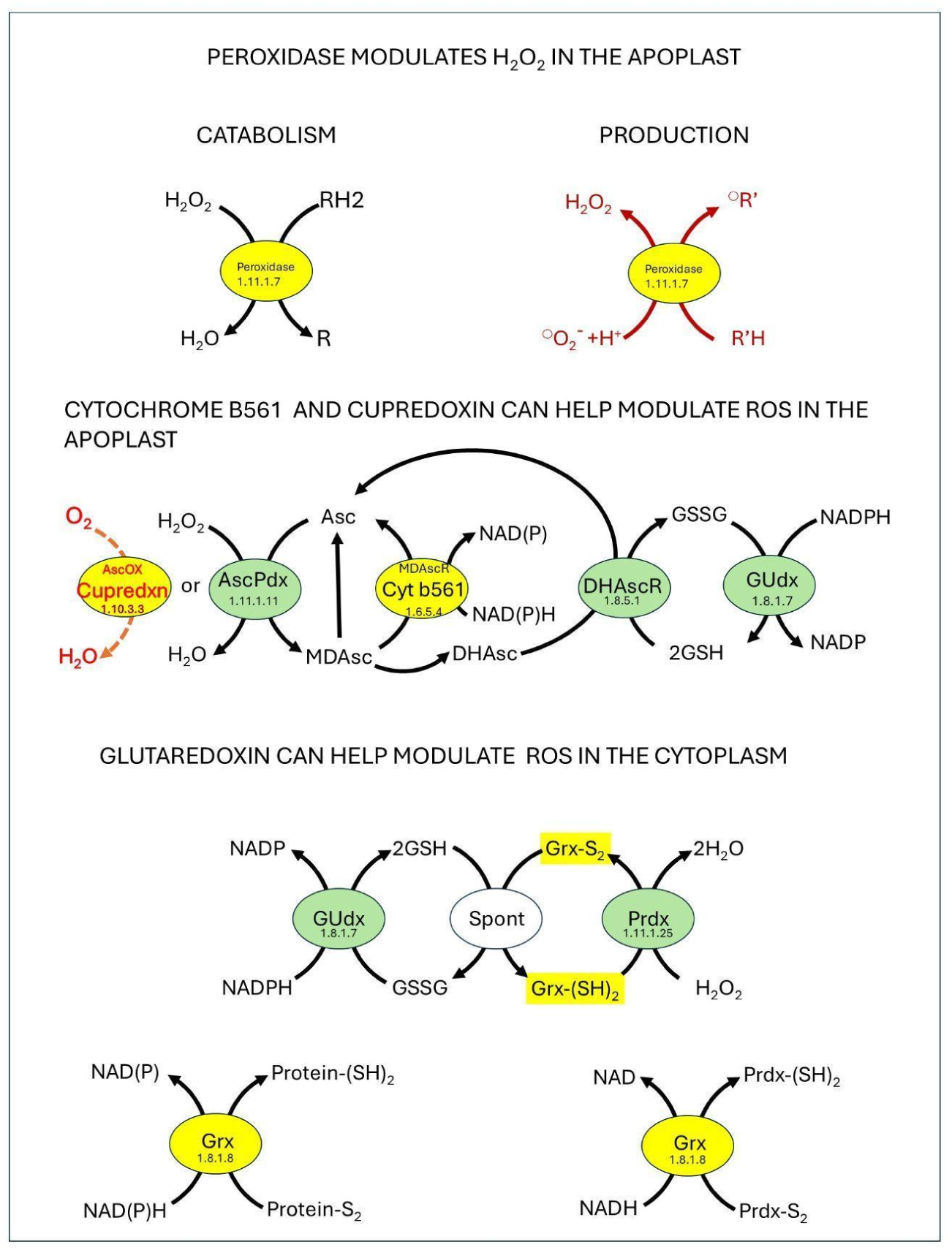
*Colletotrichum sublineola* modulates Sorghum’s oxidative stress response. The effectors are represented by yellow ovals. Top panel: double role of peroxidase in generation / degradation of hydrogen peroxide, depending on the stage of the fungal biotrophic infection. Central panel: ascorbate-glutathione cycle (<u>Do H et al., 2014</u>; <u>Kunert KJ et al., 2023</u>). Bottom panel: spontaneous reaction of glutaredoxin with glutathione (for mechanism see <u>Iversen et al., 2010</u>) and as an enzyme (EC 1.8.1.8) in the reduction of protein-disulfides. Abbreviations: **AscOx**, Ascorbate oxidase (EC 1.10.3.3); **Cupredxn**, Cupredoxin; **AscPdx**, Ascorbate peroxidase EC 1.11.1.11); **MDAscR**, Monodehydroascorbate reductase (EC 1.6.5.4); **DHAscR**, Dehydroascorbate reductase, glutathione dehydrogenase (ascorbate) (EC 1.8.5.1); **Prdx**, Peroxiredoxin (EC 1.11.1.25); **Prdx-(SH)2**, Peroxiredoxin with reduced cysteine residues; **Prdx-S2**, Peroxiredoxin with cystine disulfide bridge; **Grx**, Glutaredoxin; disulfide reductase, protein disulfide reductase (EC 1.8.1.8); **Grx-S2**, Glutaredoxin with cystine disulfide bridge; **Grx-(SH)2**, Glutaredoxin with reduced cysteine residues; **RH2,** Reduced electron donor; **R**, Oxidized electron donor; **°R’**, Substrate (Radical species form); **R’H**, Reduced substrate; **°O2-**, Superoxide; **spont**, Spontaneous; **Asc**, ascorbate; **MDAsc**, Monodehydroascorbate; **DHAsc**, Dehydroascorbate; **Protein-(SH)2**, Reduced cysteine residues in protein; **Protein-S2**, Protein with cystine disulfide bridge; **Cyt b561**, cytochrome b561. **GSH**, glutathione; **GSSG**, glutathione disulfide. **GUdx**, glutathione disulfide reductase (EC 1.8.1.7). MDAsc can spontaneously disproportionate to form ascorbate anions and water. It may also react with superoxide to form DHAsc (<u>Njus D et al., 2023</u>).

A0A066XED8 has a cytochrome b561 domain with a monodehydroascorbate reductase function. Although identified as an effector (SignalP 6.0: 0.9998; EffectorP 3.0: apoplastic 0.572), this protein has 4 transmembrane loops and 2 anchors. It plays an important role in ROS modulation, acting in concert with ascorbate and secreted fungal cupredoxins with ascorbate oxidase activity (<u>Asard H et al., 2013</u>; <u>Hu J et al., 2022</u>), like A0A066XLT0 **(Figure 4, central panel).**

*Pyricularia oryzae* MoPtep1 is a cupredoxin effector, which is targeted to the peroxisome, and modulates ROS production, but its mode of action is unknown (<u>Ning N et al., 2022</u>). As previously noted, apoplastic superoxide (^⋅^O₂⁻) is rapidly converted to H₂O₂ by superoxide dismutase. However, in the presence of transition metals such as copper, H₂O₂ can generate hydroxyl radicals via the Fenton reaction. A second role of fungal cupredoxin effectors is the mitigation of ROS generation by chelating free copper ions, thereby preventing radical formation (<u>Manzl C et al., 2004</u>; <u>Günther KL, 2018</u>). Additionally, copper release from these effectors during the necrotrophic phase may contribute to programmed cell death. Cupredoxins like *Pyricularia oryzae* MoPtep1 (<u>Ning N et al., 2022)</u>, and *Podosphaera xanthii* KX278425 (<u>Martínez-Cruz J et al., 2018</u>) **(Table 4_Oxidative_stress_Tab**), could function in this way, but this remains to be tested. Many Cs cupredoxins are not considered effectors by the SignalP 6.0/ EffectorP 3.0 criteria, however, several of them are homologs of *Podosphaera xanthii* KX278425. The effector A0A066XKN0, presents some homology to *Podosphaera xanthii* KX278425 **(Table 4 same Tab)**, which might act as a copper chelating agent, affecting Cs pathogenicity. The bacterial insect pathogen *Photorhabdus luminescens* secretes the glutaredoxin effector YdhD, which modulates oxidative stress in insect hemolymph (<u>Munch A et al., 2008</u>). Glutaredoxins catalyze the reversible reduction of disulfide bonds between glutathione and protein cysteines, maintaining redox homeostasis and participating in signaling pathways (<u>Tamayo E et al., 2016</u>). The Cs cytoplasmic glutaredoxin A0A066X0P5 may act synergistically with host enzymes such as glutathione reductase (EC 1.8.1.7), and glutaredoxin-dependent peroxiredoxin (EC 1.11.1.25), controlling H_2_O_2_ levels **(Figure 4, bottom panel).** Glutaredoxin also plays an enzymatic role (EC 1.8.1.8), using NAD(P)H + H+ to reduce disulfide bridges in other proteins, including peroxiredoxin **(Figure 4, bottom panel).**

##### Effectors Targeting ROS-Producing Systems

Some effectors disrupt host ROS production pathways. For example, *Pyricularia oryzae* AVR-Pii inhibits the NADP-malic enzyme (<u>Singh R et al., 2016</u>), blocking electron flow to NADPH oxidase. *Puccinia striiformis* effector Pst_12806 interacts with TaISP (a putative component of the cytochrome b6-f complex), likely impairing chloroplast electron transport (<u>Xu Q et al., 2019</u>). *Magnaporthe oryzae Avr*-Pita, which is an M35 metalloprotease, targets the mitochondrial OsCOX11 protein (a cytochrome c oxidase assembly protein) (<u>Han J et al., 2021</u>). Cs lacks an *Avr*-Pita homolog, but it encodes a cytoplasmic effector (A0A066XCA8) containing a cytochrome c oxidase assembly factor domain, potentially involved in regulating mitochondrial ROS. Cs also has M35 metalloprotease effectors, whose potential targets have not been identified yet (see protein degradation section).

##### Effectors Modulating ROS by Unknown Mechanisms

Certain effectors enhance fungal virulence by altering the oxidative burst through mechanisms yet to be elucidated. FgEps1, a *Fusarium graminearum* protein disulfide isomerase, induces ROS accumulation in *Nicotiana benthamiana*, though its precise function remains unclear (<u>Liu K et al., 2023</u>). Cs encodes related PDI homologs, discussed separately in the section on protein modifications **(Table 4).** Additionally, some effectors may influence ROS indirectly via vesicle trafficking, metabolite priming, or modulation of kinase signaling cascades (<u>Jwa NS and Hwang BK, 2017)</u>.

#### Network III) Proteins Modifications, Degradation and Folding

##### Protein Modifications

Pathogens use effector-mediated post-translational modifications (PTMs) to regulate infection, targeting specific proteins at defined times and locations. PTMs are especially important for biotrophic and hemibiotrophic fungi, which must suppress plant cell death during early infection. Common PTMs include glycosylation (<u>Liu C et al., 2021</u>), lipidation (myristoylation and palmitoylation), proteolysis, phosphorylation, acetylation, AMPylation, ubiquitination, and SUMOylation (<u>Tahir J et al., 2019</u>; <u>Salomon D and Orth K, 2013</u>). These modifications alter protein localization and function of the modified proteins (<u>Salomon D and Orth K, 2013</u>), helping suppress plant-triggered immunity (PTI) and enhancing virulence (<u>Tahir J et al., 2019</u>). These PTMs can occur within the pathogen, as is the case for most glycoproteins, which are glycosylated in the ER and Golgi system before being secreted by the pathogen (<u>Chen XL et al., 2020</u>). In fact, PTMs are also associated with the development of the appressorium and its penetration of the plant cells (<u>Liu C et al., 2021</u>). Many PTMs also take place within the host, where effectors modify host proteins and, in some cases, other effectors (<u>Lin B et al., 2020</u>; <u>Chen XL et al., 2014</u>). Additionally, host enzymes may modify pathogen effectors (<u>Tahir J et al., 2019</u>).

###### Glycosylation

The three principal types of protein glycosylation are N-glycosylation, O-glycosylation, and glycosylphosphatidylinositol (GPI) anchoring. Fungi utilize enzymes such as N-acetylglucosaminyl transferases, involved in N-glycan biosynthesis and ER quality control, to construct a glycosylation-based virulence network. GPI anchoring and O-mannosylation of select proteins further contribute to pathogenicity (<u>Liu C et al., 2021</u>).

###### Glycosylation’s Role in Fungal Pathogenesis

Glycosylation significantly influences protein structure and function. In some fungi, up to 72% of secreted proteins are extensively glycosylated, often at numerous sites (dozens-hundreds) (<u>Gonzalez et al., 2012</u>). Proteins linked to metabolism, cell wall formation, glycosylation, and ER quality control commonly show high N-glycosylation levels (<u>Chen XL et al., 2020</u>). Mutations in fungal N-glycosylation or GPI pathways often induce reactive oxygen species (ROS) accumulation in host tissues at infection onset, suggesting that glycosylated fungal surface proteins suppress PAMP-triggered immunity (<u>Liu C et al., 2021</u>). As previously noted, biotrophic and hemibiotrophic fungi regulate effector activity through glycosylation and deglycation. For instance, the *Magnaporthe oryzae* chitin-binding effector Slp1 [LYSM1_PYRO7] is modulated via N-glycosylation by Alg3 [M1RXP4_PYROR] (<u>Chen XL et al., 2014</u>). Similarly, O-glycosylation controls Pit1 activity, which in turn regulates the effector Pit2 (<u>Fernández-Álvarez A et al., 2012</u>).

There is substantial evidence linking fungal cell wall penetration to protein glycosylation. Successful penetration requires a structurally consistent fungal cell wall to support peg formation and turgor pressure. This integrity depends on components such as chitin, glycoproteins, and β-1,3-glucans and is influenced by glycosylation or deglycation of wall proteins (<u>Lesage G and Bussey H, 2006</u>; <u>Samalova M et al., 2017</u>) and GPI anchoring, as demonstrated in *M. oryzae* (<u>Fujita M and Kinoshita T, 2012</u>).

###### Glycosylation’s Indirect Role in Appressorium Penetration

Protein glycosylation can influence appressorium function through indirect mechanisms. For instance, deletion of pmt4, an O-mannosyltransferase, disrupts appressorium formation and host penetration by impairing downstream activation of the MAP kinase Kpp2. Similarly, in *M. oryzae*, N-glycosylation of ALG3 affects appressorium functionality (<u>Liu C et al., 2021</u>). **Table 4 (Glycosylation_Tab)** outlines examples of glycosylation’s role in fungal pathogenicity.

###### Modifications Involving Glycosyl Hydrolases and Effector Proteins

Glycosyl hydrolases (GHs), a subclass of CAZymes, catalyze the cleavage and restructuring of glycosidic bonds in polysaccharides and glycoconjugates. With over 170 families, many possessing diverse substrate specificities, hemibiotrophic and necrotrophic fungi typically encode around 300 GH genes. Their GH repertoire often reflects the host plant’s cell wall composition, which differs between monocots and dicots (<u>Bradley EL et al., 2022</u>). Effector proteins must be correctly folded and functional for successful infection. The <u>E</u>ndoplasmic <u>R</u>eticulum <u>Q</u>uality <u>C</u>ontrol (ERQC) system ensures that only properly structured glycoproteins are secreted. Accordingly, glucosidase I enzymes that remove glycosyl residues from misfolded proteins in the fungal ER are critical for pathogenicity (<u>Liu C et al., 2021</u>).

Conversely, some glycosylated effectors are secreted via the classical pathway and subsequently activated in the host by partner effectors with GH activity. Early in infection, fungi release GHs and other lytic enzymes to degrade host cell walls and facilitate invasion. Usually, glycoside hydrolases (GHs) serve two functions: (1) they facilitate appressorium penetration by disrupting the plant cell wall, and (2) they provide nutrients (cell wall degradation products). Some of these products also act as elicitors, triggering immune responses that may lead to host cell death (<u>Wan J et al., 2021</u>). Hemibiotrophic fungi counteract this by employing mechanisms early in infection to suppress such responses, as discussed above in the “Apoplast Immune Mechanisms Interplay” section. Additionally, certain secreted GHs have a second effector role, beyond that of cell wall degradation alone, since they remove glycosyl moieties from specific proteins to enhance virulence. Several GH families with this activity have been identified (**Table 4_GH_other_roles_Tab)**. GHs may also detoxify plant-derived compounds or modulate immune responses (<u>Bradley EL et al., 2022</u>).

Cs possesses several GH effector candidates, classified into superfamilies GH7, GH10, GH11, GH12, GH16, GH28, GH35, GH43, GH75, and GH131 **(Table 2_Glycosyl_hydrolases_Tab)**.

Most are predicted to localize exclusively to the apoplast, except A0A066WZH1 (GH131) and A0A066X4Y3 (GH10), which are cytoplasmic, and A0A066XGR6 (GH11), which exhibits dual localization. Based on domain architecture and apoplastic localization, these GHs likely perform CAZyme-associated enzymatic functions **(Table 2_Glycosyl_hydrolases_Tab).** Notably, some GHs—particularly those from families GH11, and GH28, share high sequence similarity with experimentally validated fungal effectors known to enhance virulence **(Figure 5).** This supports their classification as bona fide Cs effectors by SignalP 6.0 and EffectorP 3.0 and suggests additional roles in deglycosylation and virulence modulation. For instance, A0A066X6S5 is homologous to *Botryotinia fuckeliana* endo-polygalacturonase Q9Y7V9, which is recognized by a leucine-rich repeat receptor-like protein (RLP PRR) and modulates host cell death. Similarly, endo-1,4-β-xylanase P0CT49 regulates virulence in *Pyricularia oryzae*; its closest Cs homolog is A0A066XFB3, though homologs A0A066XLZ4 and A0A066X9Z0 also show similarity, leaving their functional equivalence to P0CT49 unresolved.

**Figure 5.**
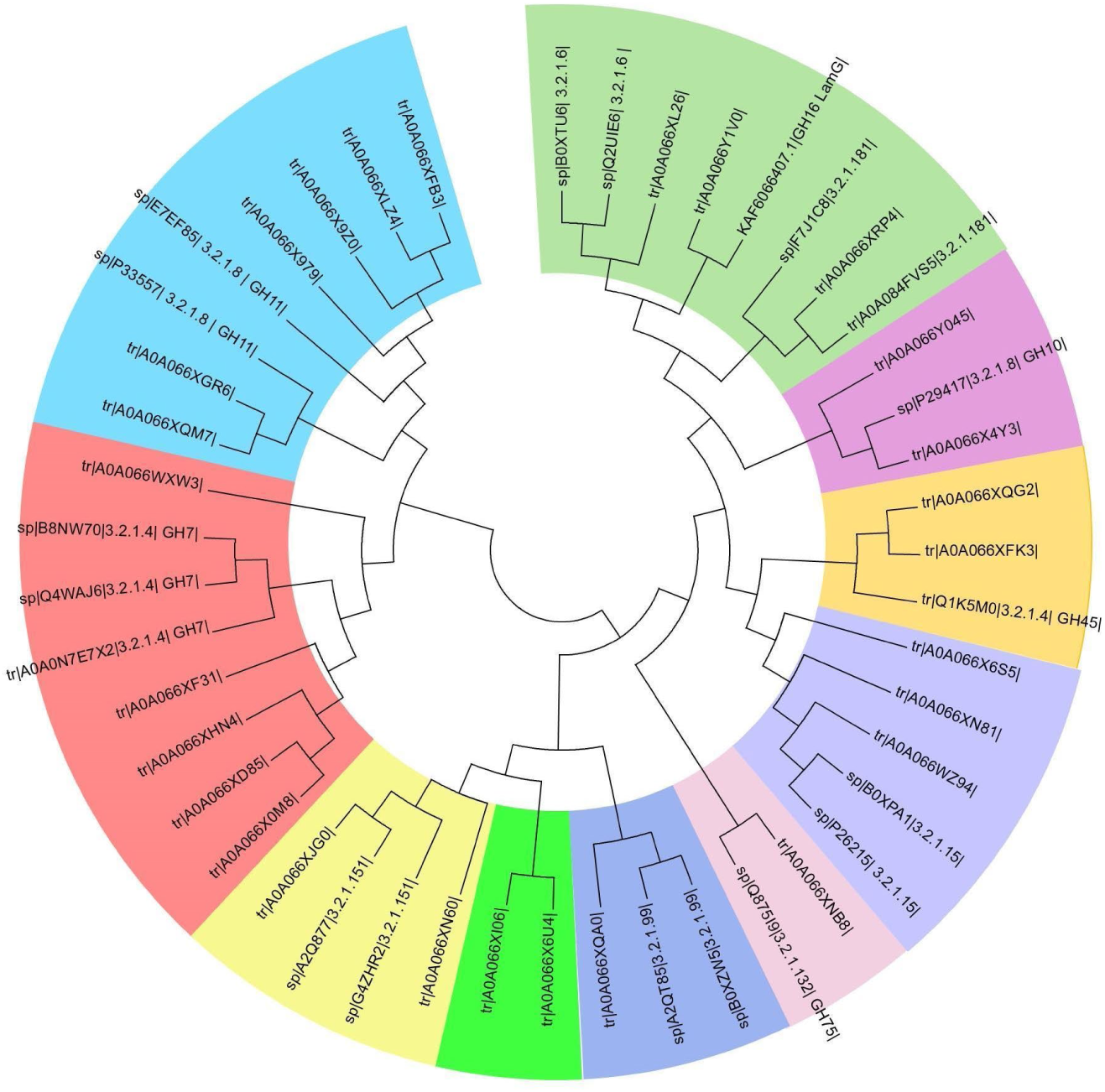
Phylogenetic tree of *Colletotrichum sublineola* glycosyl hydrolase effectors. Evolutionary tree made using the Maximum Likelihood method conducted with the MEGA6 program. The bootstrap consensus tree represents the evolutionary history of the Cs glycosyl hydrolase effectors, and this tree is inferred from 100 replicates. Cs has 28 effectors with glycosyl hydrolase domains (GH 7, 10,11,12,16,28,43,45,75, and 131). For details see **Table 2**. We excluded A0A066WZH1 because it has 2 different GH domains. Sequences from known glycosyl hydrolases were obtained from the BRENDA and UniProt databases and added as markers. Some GHs—particularly those from families GH11 (EC 3.2.1.8; baby blue clade), and GH28 (EC 3.2.1.15; purple clade), share high sequence similarity with experimentally validated fungal effectors known to enhance virulence.

##### Protein Isomerization

###### Protein Disulfide Isomerases

Protein disulfide isomerases (PDIs) catalyze disulfide bond formation, breakage, and rearrangement across various proteins. While primarily involved in ER-associated protein folding, PDIs also function in other cellular compartments, including roles in NADP(H) oxidase regulation and microsomal triglyceride transfer <u>(Hatahet F & Ruddock LW, 2009</u>). In pathogens such as *Leishmania major* and *Entamoeba histolytica*, PDIs modulate infection mechanisms by influencing ROS production by NADPH oxidase, antigen processing in the ER (protein folding), and host invasion (<u>Stolf BS et al., 2011</u>). Cs encodes two PDI effectors from superfamily cl36828 (A0A066X7Y3 and A0A066X8J7) and two from a_ERp38; cd02998 superfamily (A0A066XAT9 and A0A066XBK1). A0A066XBK1 shares 50.7% identity (E=0) with the *Fusarium graminearum* effector FgEps1, whose deletion reduces DON toxin production and pathogenicity (<u>Liu K et al., 2023</u>), despite lacking a nuclear targeting signal. A0A066X8J7 is homologous to *Pyricularia oryzae* MoPdi1 (76.8% identity; E=0), which regulates ER stress, contributes to AVR-Pia effector secretion, and affects virulence, cell wall integrity, and conidiation **(Table 4_Protein_Modifications_Tab)**. Both A0A066X8J7 and MoPdi1 lack a predicted nuclear target peptide.

###### Peptidyl-Prolyl Cis-Trans Isomerases

Peptidyl-prolyl cis-trans isomerases (PPTIs; EC 5.2.1.8) catalyze the isomerization of proline residues in proteins (<u>Pemberton TJ, 2006</u>), contributing to protein folding and assembly, signal transduction, and cell cycle regulation (<u>Göthel SF & Marahiel MA, 1999</u>). PPTIs comprise cyclophilins, parvulins, and FK506-binding proteins. Cs encodes two PPTI effectors: A0A066X9Q0 (a cyclophilin) and A0A066XK85 (an FK506-binding protein). The *Magnaporthe grisea* cyclophilin CYP1, a virulence factor involved in appressorium function, shares 58.8% identity with A0A066X9Q0 (E = 3.7e−57), though non-effector Cs cyclophilins display even higher homology. A0A066X9Q0 also aligns well with two *Plenodomus lingam* PPTI effectors (E4ZHT6, E4ZIY5), while A0A066XK85 shares 61.1% identity with E4ZTE6 (E = 6.8e−48) **(Table 4_Protein_Modifications_Tab).** Notably, E4ZTE6 (LmFKB2) exhibits low expression during infection, consistent with the pathogen’s transition from saprophytic to hemibiotrophic growth (<u>Singh K et al., 2014</u>).

###### Protein Degradation

Metalloproteases (MPs) contain a divalent metal cation in their catalytic site. Most exhibit an HEXXH+D sequence (the aspzincin motif), where two histidine residues coordinate zinc binding. Among pathogenic fungi, the major secreted MPs include deuterolysins (M35) and fungalysins (M36); the latter also possess a C-terminal GTXDXXYG motif, with an aspartate residue completing zinc coordination (<u>Pan L et al., 2020</u>). Pappalysins (M43), shorter in fungi and bacteria than in vertebrates, consist of a catalytic and a pro-domain and harbor the HEXXHXXGXXH zinc binding motif (<u>Courrol DDS et al., 2022</u>).

Cs encodes two M43 apoplastic effectors (A0A066X544, A0A066XE02) and four M35 effectors—one apoplastic (A0A066XJ51) and three cytoplasmic (A0A066XH92, A0A066XHE8, A0A066XRK6). A0A066X544 and A0A066XE02 share high similarity with *Rhizoctonia cerealis* RcMEP1 (44.9% and 43.4% identity; E = 4.9e−72 and 8.3e−64, respectively) **(Table 4_Protein_Modifications_Tab).** RcMEP1 is a metalloprotease that induces ROS accumulation, cell death, and suppresses host chitinase accumulation (<u>Pan L et al., 2020</u>). A0A066XRK6 is homologous to *Colletotrichum higginsianum* A0A1B7YCE5_COLHI, though its function remains uncharacterized. A0A066XJ51 also shows moderate homology to *Verticillium dahliae* VdM35-1 (27.7% identity; E = 5.6e−24), an effector that activates ROS production and host cell death (<u>Lv J et al., 2022</u>).

Additionally, Cs encodes seven apoplastic effectors from the α/β-hydrolase superfamily (cl21494). These enzymes, which share a nucleophile–His–acid catalytic triad, act on diverse substrates (<u>Holmquist M, 2000</u>). For instance, A0A066XPB8 is homologous to *Sclerotinia sclerotiorum* SsCut1 (61% identity; E = 1.3e−93), a hydrolase with cutinase activity (<u>Gong Y et al., 2022</u>).

##### Protein Folding and Candidate Effectors that Could Hijack it

During pathogen attack plants experience ER stress leading to the accumulation of misfolded or unfolded proteins, triggering the unfolded protein response (UPR). If unresolved, UPR may culminate in cell death. ER protein folding occurs via N-glycan–independent and N-glycan–dependent pathways. In the former, ERdj3 binds to nascent proteins in the reticulum, and proceeds to bind BiP, which hydrolyzes ATP. The BiP-ADP chaperone then binds proteins with high affinity. Finally, an exchange of ATP for the bound ADP frees the protein, which continues folding (<u>Deng Y et al, 2013</u>). The ATPase cycle of the Bip is regulated by the Sil1 and Grp170 (glucose-regulated protein) (<u>Behnke J et al, 2015</u>). In the N-glycan–dependent pathway, glucosidases I and II, lectin chaperones (calnexin and calreticulin), and foldases promote proper protein folding (<u>Deng Y et al., 2013</u>). Below, we examine effectors containing domains associated with these chaperones and assess their potential involvement in plant ER folding mechanisms.

###### Calreticulins

The Cs cytoplasmic effector A0A066X6Y7 contains a calreticulin domain. While Localizer does not identify a transit peptide, WoLFPSORT suggests extracellular and ER/Golgi localization in plants, implying a potential role in host protein folding. Given calreticulin’s diverse functions beyond chaperoning (<u>Nakhasi HL et al., 1998</u>), subcellular localization is critical. Notably, calreticulin effectors from *Radopholus similis* and *Meloidogyne incognita* suppress plant defenses and possess nuclear transit peptides (<u>Jaouannet M et al., 2013</u>; <u>Li Y et al., 2015</u>) **(Table 4_Protein_Modifications_Tab)**. A0A066X6Y7 shows strong homology to both *R. similis* AFK76483 (37.4 % with E=3.9 e-80) and *M. incognita* Mi-CRT (37.3 % and E=1.4 e-81) effectors **(Table 4, same Tab).**

###### Glucosidase II

A0A066XM65 is a Cs effector featuring a central ATPase domain flanked by two glucosidase II domains, which are involved in recognizing misfolded glycoproteins. WoLFPSORT predicts ER localization, whereas Localizer identifies a nuclear transit peptide in A0A066XM65. This protein shares some homology with the *Mycosarcoma maydis* Gas2 effector (27.3 % with E=2.6 e-61), which facilitates fungal spread within host tissues **(Table 4, same Tab**), however, there are some differences in the central domain composition.

###### DnaJ/Hsp40 Protein Effectors

DnaJ proteins (Hsp40 family) exhibit diverse domain architectures and are classified into three types:

-Type I: Contains an N-terminal J domain (interacts with Hsp70 and has ATPase activity), a glycine/phenylalanine-rich (G/F) region, and a cysteine-rich zinc-binding domain.

-Type II: Possesses the J and G/F regions but lacks the cysteine-rich domain.

- Type III: Contains only the J domain, with a variety of locations within the protein (Cheetham ME and Caplan AJ, 1998).

The Cs genome encodes three DnaJ/Hsp40 effectors: one shows medium homology and two very low homology to the DnaJ effectors which have been studied in other organisms **(Table 4_Protein_Modifications_Tab).** These effectors are:

- A0A066XBI3: A canonical type I DnaJ protein. In *Escherichia coli* and *Clostridium difficile*, DnaK is involved in heat tolerance (Ogura M et al., 2025). The bacterial Dnaj chaperone is normally part of the DnaK-DnaJ-GrpE system, and when the Hsp70 ATPase is stimulated, this system prevents unfolded proteins to assemble (NCBI). A0A066XBI3 presents some homology to the *Pseudomonas cichorii* JBC1 DnaJ effector (33.3% E= 2.6 e-45) **(Table 4, same Tab),** which is linked to oxidative stress response. Probably A0A066XBI3 may similarly regulate some stress responses during the Cs infection process.

- A0A066WYT5: A type III DnaJ protein with a predicted nuclear transit peptide. It contains an N-terminal J domain but lacks the G/F and cysteine-rich regions. A central ribonuclease domain **(Table 1)** suggests a potential role in processing host RNA precursors.

- A0A066X1N3: Features a C-terminal J domain followed by G/F region and a lipopolysaccharide biosynthesis regulator YciM/LapB domain near the N terminal **(Table 1).** This domain includes six tetratricopeptide repeats and a C-terminal metal-binding motif. In bacteria, LapB by itself normally modulates cellular lipopolysaccharide levels. The functional implications of this DnaJ–LapB fusion in the effector remain unclear.

###### Hsp70 and SIL1 Effectors

Hsp70 proteins are multifunctional chaperones involved in protein folding, signal transduction, and translocation across various cellular compartments (<u>Xu L et al., 2018</u>). A0A066WXV8 is the only identified Cs effector containing an Hsp70 domain and is predicted to localize to the plant nucleus (KKNNVDITKDLKAMGKLKR,KKTMKPVEQVLKDAKIKKE) according to Localizer (https://localizer.csiro.au/). As no Hsp70 effectors have been reported to date, this nuclear targeting—if not a false positive—suggests a function distinct from ER-associated protein folding. Notably, some Hsp70 proteins can bind DNA (<u>Zeiner M et al., 1999</u>), which may also apply to A0A066WXV8. Also, A0A066X8N1 encodes a SIL1-like nucleotide exchange factor, known to interact with Hsp70 chaperones (<u>Behnke J et al., 2015</u>). However, Localizer does not predict a nuclear localization signal for this protein.

#### Network IV) CFEM Fungal Proteins and Modulation of Plant Immunity

CFEM domains are cysteine-rich motifs unique to fungi (<u>Kulkarni RD et al., 2003</u>). CFEM has the consensus sequence <PxC[A/G]x₂Cx₈₋₁₂Cx₁₋₃[x/T]Dx₂₋₅CxCx₉₋₁₄Cx₃₋₄Cx₁₅₋₁₆>. While many fungal genomes encode CFEM proteins, not all are secreted effectors (many are membrane-bound). Not all fungi have CFEM proteins and pathogenic fungi typically possess more CFEM proteins than other fungi (<u>Zhang ZN, et al., 2015</u>).

Some CFEM effectors modulate plant immunity by interacting with host immune regulators and mediators altering ROS production and programmed cell death. Others influence appressorium development and structure **(Table 4_CFEM_Tab).** In the Cs genome, 18 CFEM proteins were identified **(Table 2),** including five predicted apoplastic effectors: A0A066X314, A0A066X590, A0A066X5K5, A0A066XGJ6, and A0A066XK30, whereas A0A066XD31 is predicted to be dual-targeted.

Structural conservation is evident among CFEM effectors in the *Colletotrichum* genus **(Figure 6).** For example, Cs effector A0A066XD31 shares 98% identity with *Colletotrichum graminicola* CgCFEM8 (E3Q7L5) (E = 1.1 e-70), while A0A066XGJ6 is 81.4% identical to CgCFEM9 (E3QD10) (E = 8.6 e-84) **(Figure 6).** Both CgCFEM8 and CgCFEM9 are proposed to suppress hypersensitive response (HR) in host plants (<u>Gong AD et al., 2020</u>). A0A066XD31 and CgCFEM8 also show homology to *Colletotrichum fruticola* CfEC12, **(Figure 6).** Plants’ NIMIN (NIM1-interacting) proteins interact with the regulatory factor NPR1(<u>N</u>on-expressor of <u>P</u>athogenesis-<u>R</u>elated genes 1) to control the expression of defense genes (<u>Zavaliev R and Dong X, 2024</u>) NPR1 itself is a receptor of salicylic acid. The CfEC12 effector competes with the host’s NPR1 for a target binding site in NIMIN2 and suppresses the interaction between NIMIN2 and NPR1, thus attacking the plant’s salicylic acid defense pathway (<u>Shang S et al., 2024</u>). Given its dual apoplastic and cytoplasmic localization, A0A066XD31 may similarly interfere with immune regulators in both compartments, potentially targeting the salicylic acid defense pathway in *Sorghum*. This hypothesis remains to be experimentally validated. Additionally, two studies of CFEM proteins’ role in *Magnaporthe oryzae* (also known as *M. grisea* and *Pyricularia oryzae*) reveal that MoCDIP2 (also identified as AAN64312.1) is associated with host cell death modulation and appressorium differentiation. This effector is a good homolog of A0A066X5K5 (43.2% identity, E = 1.1e-38) **(Table 4_CFEM_Tab).**

**Figure 6.**
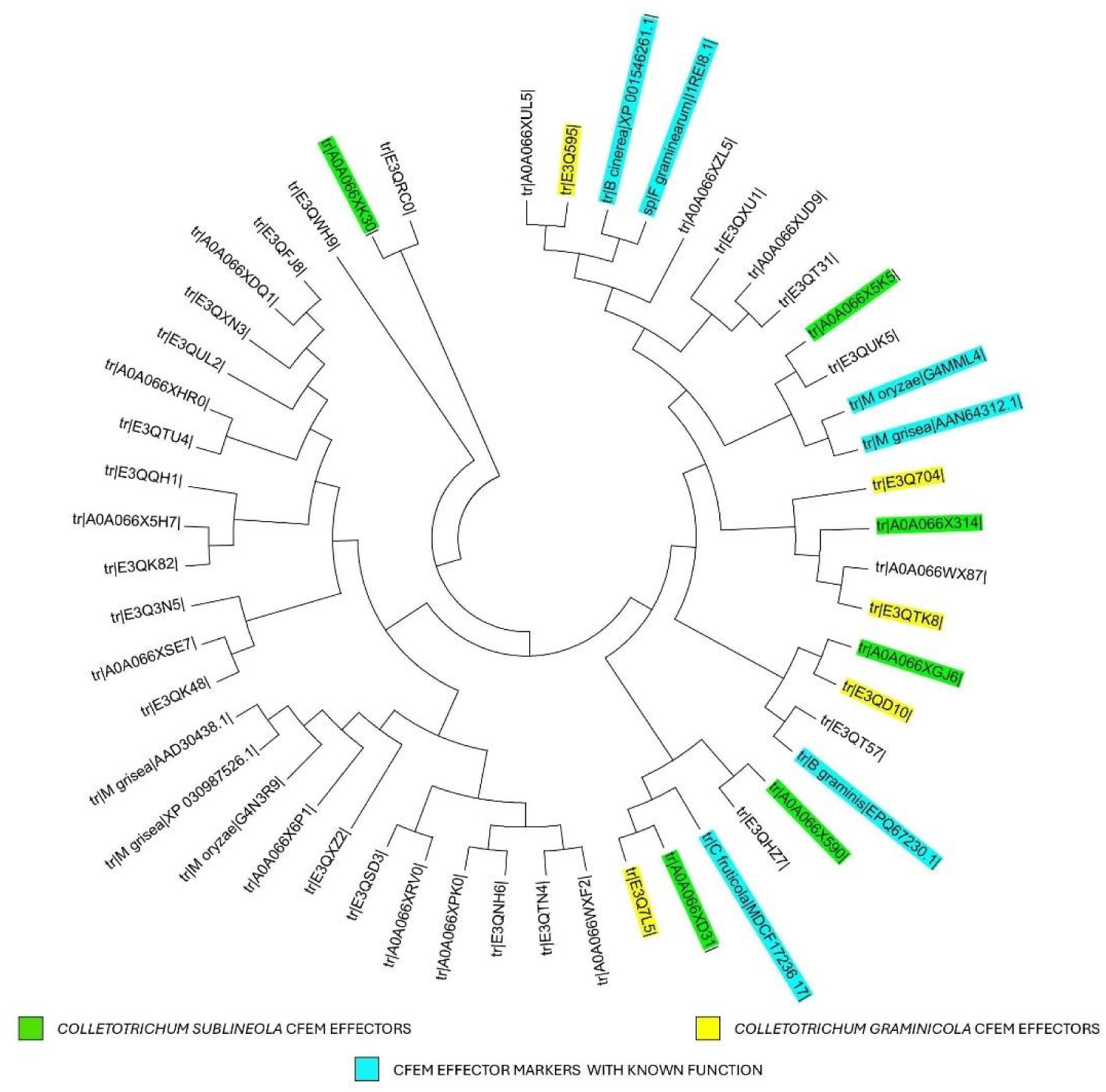
Phylogenetic tree of the *Colletotrichum sublineola* CFEM protein family. Evolutionary tree made using the Maximum Likelihood method conducted with the MEGA6 program. The bootstrap consensus tree represents the evolutionary history of the CFEM proteins and is inferred from 100 replicates. Originally 51 amino acid sequences were involved, but the positions containing gaps and missing data were eliminated. The final dataset contained 42 positions. Only some members of the CFEM family are effectors, and other membranes are membrane proteins **(Table_2_CFEM_Tab**). The effectors are color tagged and some fungal CFEM effectors with known functions were added as markers (see above).

## Discussion

Plant pathogenic fungal expression data are difficult to obtain because the amount of RNA extracted from samples at different stages of invasion is small compared to the host’s own RNA (<u>Vela S et al., 2024</u>). Despite this, these data support the hypothesis that effectors’ expression profiles change during infection stages; however, it remains unclear whether we are capturing the complete spectrum of these effectors in this type of analysis. Conversely, a comprehensive analysis of the effectors present within a fungal proteome offers valuable insights into its pathogenic potential. This analysis starts with genome sequencing and assembly and is followed by computational analyses that identify putative effectors. Despite their utility, these methods carry their own inherent limitations and definitive characterization of each effector is required for accurate determination of their function. Given the abundance of potential effector proteins, experimental validation of all candidates remains challenging. In light of these constraints, we adopted computational approaches - notably comparative genomics- to predict the functions of several putative effectors of Cs, which were identified using SignalP 6.0 and EffectorP 3.0. Nearly half of the Cs proteins identified as putative effectors have conserved domains, highlighting the benefit of applying these predictive methods to prioritize the most promising candidates for subsequent experimental validation.

Due to horizontal gene transfer between plant pathogens (<u>Mehrabi R et al., 2011</u>), we were able to extend even further our comparative genomics approach. This was based on studies done not only in fungi, but also in organisms as different as nematodes (<u>Haegeman A et al., 2011</u>) **(Table 4_Protein Modifications_Tab).** We organized the predicted functions of Cs effectors into subsystems, which enabled us to integrate isolated functions into more clearly defined pathogenic mechanisms. However, as we have observed, comparative genomics also has its limitations. For example, proteins that serve as effectors in some fungi can share high sequence homology with proteins from other fungi that are not effectors **(see Table 4_CFEM_Tab).** Furthermore, effectors roles are complex and can be affected by moonlighting proteins, which perform multiple, unrelated functions within the same organism (<u>Arvizu-Rubio VJ., et al, 2022</u>). Additionally, while some effectors may maintain their enzymatic activities, they might target similar proteins in different systems, leading to varied effects **(Table 4_GH_Tab).** Some of the predicted Cs effectors with conserved domains have remained unexplored and so far, we have only classified them into the following small groups: inhibitors, virulence factors, signaling proteins, allergens, and necrosis-inducing factors **(Table 1).** Predicting their functions remains a challenge for future research. Finally, in the case of effectors with non-conserved domains, their functional specificity is even greater, and very few have sequence homologs in other fungi, let alone sequence homologs showing functional role similarity, so comparative genomics analysis is not applicable in their case. Although effectors with non-conserved domains exhibit high sequence variations, some conserve tertiary structure motifs which may be associated with similar functions (<u>Rozano L et al., 2024</u>). In this case, the structural analysis provided by advanced computational methods like AlphaFold has been invaluable (<u>Seong K et al., 2023</u>).

In this study we identified candidate effectors in Cs and used comparative genomics to predict their functions. This work may inform the development of targeted strategies to mitigate anthracnose. For example, cepharanthine (CEP), a compound known to bind the FsCFEM1protein, has been shown to inhibit growth in multiple *Fusarium* species (<u>Wang Y et al., 2025</u>). Our analysis reveals the presence of several CFEM-domain-containing proteins within the Cs effector repertoire. Although none of the Cs CFEM proteins is a homolog to this *Fusarium* CFEM motif, the results with CEP warrant investigation of chemicals that by reacting with CFEM groups could work as therapeutic agents against anthracnose. Another example is that of *Alternaria alternata* mutants lacking the Glr1 glutaredoxin, which exhibit heightened sensitivity to the fungicide mancozeb (<u>Ma H et al., 2018</u>). Given that Cs genome encodes three glutaredoxins, one classified as an effector, targeting of Cs glutaredoxins, may enhance susceptibility to mancozeb or structurally related fungicides. This is an example of the applications that could be brought by the discovery of Cs effectors functions, encouraging researchers to continue exploring them.

## Tables

This publication tables can be viewed and downloaded from this figshare webpage: https://doi.org/10.6084/m9.figshare.31347355

## Acknowledgements

This work is supported as part of the BER BRaVE project KP1601011 and by KBase. KBase and BRaVE are funded by the U.S. Department of Energy, Office of Science, Office of Biological and Environmental Research under Award Numbers DE-AC02-06CH11357, and DE-AC02-98CH10886.

## Acknowledgements

The submitted manuscript has been created by UChicago Argonne, LLC, Operator of Argonne National Laboratory (“Argonne”). Argonne, a U.S. Department of Energy Office of Science laboratory, is operated under Contract No. DE-AC02-06CH11357. The U.S. Government retains for itself, and others acting on its behalf, a paid-up nonexclusive, irrevocable worldwide license in said article to reproduce, prepare derivative works, distribute copies to the public, and perform publicly and display publicly, by or on behalf of the Government. The Department of Energy will provide public access to these results of federally sponsored research in accordance with the DOE Public Access Plan. http://energy.gov/downloads/doe-public-access-plan

## References

Abreha KB, Ortiz R, Carlsson AS, Geleta M (2021) Understanding the Sorghum-Colletotrichum sublineola Interactions for Enhanced Host Resistance. Front Plant Sci.12:641969. doi: 10.3389/fpls.2021.641969.

Arvizu-Rubio VJ, García-Carnero LC, Mora-Montes HM (2022) Moonlighting proteins in medically relevant fungi. PeerJ.10:e14001. doi: 10.7717/peerj.14001.

Asard H, Barbaro R, Trost P, Bérczi A (2013) Cytochromes b561: ascorbate-mediated trans-membrane electron transport. Antioxid Redox Signal.19:1026–1035. doi: 10.1089/ars.2012.5065.

Bateman A, Bycroft M (2000) The structure of a LysM domain from E. coli membrane-bound lytic murein transglycosylase D (MltD). J Mol Biol. 299:1113–1119. doi: 10.1006/jmbi.2000.3778.

Behnke J, Feige MJ, Hendershot LM (2015) BiP and its nucleotide exchange factors Grp170 and Sil1: mechanisms of action and biological functions. J Mol Biol. 427:1589–1608. doi: 10.1016/j.jmb.2015.02.011.

Bradley EL, Ökmen B, Doehlemann G, Henrissat B, Bradshaw RE, Mesarich CH (2022) Secreted Glycoside Hydrolase Proteins as Effectors and Invasion Patterns of Plant-Associated Fungi and Oomycetes. Front Plant Sci. 13:853106. doi: 10.3389/fpls.2022.853106.

Černý M, Habánová H, Berka M, Luklová M, Brzobohatý B (2018) Hydrogen Peroxide: Its Role in Plant Biology and Crosstalk with Signalling Networks. Int J Mol Sci. 19:2812. doi: 10.3390/ijms19092812.

Cheetham ME and Caplan AJ (1998) Structure, function and evolution of DnaJ: conservation and adaptation of chaperone function. Cell Stress Chaperones 3:28–36. doi: 10.1379/1466-1268(1998)003<0028:sfaeod>2.3.co;2.

Chen XL, Shi T, Yang J, Shi W, Gao X, Chen D, Xu X, Xu JR, Talbot NJ, Peng YL (2014) N-glycosylation of effector proteins by an α-1,3-mannosyltransferase is required for the rice blast fungus to evade host innate immunity. Plant Cell. 26:1360–1376. doi: 10.1105/tpc.114.123588.

Chen XL, Liu C, Tang B, Ren Z, Wang GL, Liu W (2020) Quantitative proteomics analysis reveals important roles of N-glycosylation on ER quality control system for development and pathogenesis in Magnaporthe oryzae. PLoS Pathog. 16:e1008355. doi: 10.1371/journal.ppat.1008355.

Courrol DDS, da Silva CCF, Prado LG, Chura-Chambi RM, Morganti L, de Souza GO, Heinemann MB, Isaac L, Conte FP, Portaro FCV, Rodrigues-da-Silva RN, Barbosa AS (2022) Leptolysin, a Leptospira secreted metalloprotease of the pappalysin family with broad-spectrum activity. Front Cell Infect Microbiol. 12:966370. doi: 10.3389/fcimb.2022.966370.

De Gara L, Locato V, Dipierro S, de Pinto MC (2010) Redox homeostasis in plants. The challenge of living with endogenous oxygen production. Respir Physiol Neurobiol. 173 Suppl:S13–19. doi: 10.1016/j.resp.2010.02.007.

Demidchik V (2015) Mechanisms of oxidative stress in plants: From classical chemistry to cell biology. Environmental and Experimental Botany 109:212–228. 10.1016/j.envexpbot.2014.06.021

Deng Y, Srivastava R, Howell SH (2013) Endoplasmic reticulum (ER) stress response and its physiological roles in plants. Int J Mol Sci. 14:8188–8212. doi: 10.3390/ijms14048188.

Derbyshire MC, Raffaele S (2023) Surface frustration re-patterning underlies the structural landscape and evolvability of fungal orphan candidate effectors. Nat Commun14:5244. doi: 10.1038/s41467-023-40949-9.

De Wit PJGM, Testa AC, Oliver RP (2016) Fungal Plant Pathogenesis Mediated by Effectors. Microbiol Spectr. 4:10.1128/microbiolspec.funk-0021-2016.4(6). doi: 10.1128/microbiolspec.FUNK-0021-2016.

Do H, Kim IS, Kim YS, Shin SY, Kim JJ, Mok JE, Park SI, Wi AR, Park H, Lee JH, Yoon HS, Kim HW (2014) Purification, characterization and preliminary X-ray crystallographic studies of monodehydroascorbate reductase from Oryza sativa L. japonica. Acta Crystallogr F Struct Biol Commun. 70:1244–1248. doi: 10.1107/S2053230X14015908.

Domazakis E, Lin X, Aguilera-Galvez C, Wouters D, Bijsterbosch G, Wolters PJ, Vleeshouwers VG.(2017) Effectoromics-Based Identification of Cell Surface Receptors in Potato. Methods Mol Biol. 1578:337–353. doi: 10.1007/978-1-4939-6859-6_29

Dunyak BM, Gestwicki JE (2016) Peptidyl-Proline Isomerases (PPIases): Targets for Natural Products and Natural Product-Inspired Compounds. J Med Chem. 59:9622–9644. doi: 10.1021/acs.jmedchem.6b00411.

Fernández-Álvarez A, Marín-Menguiano M, Lanver D, Jiménez-Martín A, Elías-Villalobos A, Pérez-Pulido AJ, Kahmann R, Ibeas JI (2012) Identification of O-mannosylated virulence factors in Ustilago maydis. PLoS Pathog. 8:e1002563. doi: 10.1371/journal.ppat.1002563.

Fiorin GL, Sanchéz-Vallet A, Thomazella DPT, do Prado PFV, do Nascimento LC, Figueira AVO, Thomma BPHJ, Pereira GAG, Teixeira PJPL (2018) Suppression of Plant Immunity by Fungal Chitinase-like Effectors. Curr Biol. 28:3023–3030.e5. doi: 10.1016/j.cub.2018.07.055.

Fujita M, Kinoshita T (2012) GPI-anchor remodeling: potential functions of GPI-anchors in intracellular trafficking and membrane dynamics. Biochim Biophys Acta 1821:1050–1058. doi: 10.1016/j.bbalip.2012.01.004.

Gaderer R, Bonazza K, Seidl-Seiboth V (2014) Cerato-platanins: a fungal protein family with intriguing properties and application potential. Appl Microbiol Biotechnol. 98:4795–4803. doi: 10.1007/s00253-014-5690-y.

Gong AD, Jing ZY, Zhang K, Tan QQ, Wang GL, Liu WD (2020) Bioinformatic analysis and functional characterization of the CFEM proteins in maize anthracnose fungus Colletotrichum graminicola. Journal of Integrative Agriculture19: 541–550; 10.1016/S2095-3119(19)62675-4

Gong Y, Fu Y, Xie J, Li B, Chen T, Lin Y, Chen W, Jiang D, Cheng J (2022) Sclerotinia sclerotiorum SsCut1 Modulates Virulence and Cutinase Activity. J Fungi (Basel) 8:526. doi: 10.3390/jof8050526.

González M, Brito N, González C (2012) High abundance of Serine/Threonine-rich regions predicted to be hyper-O-glycosylated in the secretory proteins coded by eight fungal genomes. BMC Microbiol. 12:213. doi: 10.1186/1471-2180-12-213.

Göthel SF, Marahiel MA (1999) Peptidyl-prolyl cis-trans isomerases, a superfamily of ubiquitous folding catalysts. Cell Mol Life Sci. 55:423–436. doi: 10.1007/s000180050299.

Gronenborn AM (2009). Structures of Cvnh Family Lectins. In: Puglisi, J.D. (eds) Biophysics and the Challenges of Emerging Threats. NATO Science for Peace and Security Series B: Physics and Biophysics. Springer, Dordrecht. 10.1007/978-90-481-2368-1_3

Günther KL (2018) Investigation of the reactive oxygen species metabolism during the life cycle of *Fusarium graminearum*. PhD thesis, Faculty of Mathematics, Informatics, and Natural Sciences, Dept. of Biology, University of Hamburg, Germany. <u>Dissertation.</u>

Haegeman A, Jones JT, Danchin EG (2011) Horizontal gene transfer in nematodes: a catalyst for plant parasitism? Mol Plant Microbe Interact. 24:879–887. doi: 10.1094/MPMI-03-11-0055.

Han J, Wang X, Wang F, Zhao Z, Li G, Zhu X, Su J, Chen L (2021) The Fungal Effector Avr-Pita Suppresses Innate Immunity by Increasing COX Activity in Rice Mitochondria. Rice (N Y) 14:12. doi: 10.1186/s12284-021-00453-4.

Hasanuzzaman M, Fujita M (2022) Plant Oxidative Stress: Biology, Physiology and Mitigation. Plants (Basel). 11:1185. doi: 10.3390/plants11091185.

Hatahet F, Ruddock LW (2009) Protein disulfide isomerase: a critical evaluation of its function in disulfide bond formation. Antioxid Redox Signal. 11:2807–2850. doi: 10.1089/ars.2009.2466.

He Q, McLellan H, Boevink PC, Birch PRJ (2020) All Roads Lead to Susceptibility: The Many Modes of Action of Fungal and Oomycete Intracellular Effectors. Plant Commun. 1:100050. doi: 10.1016/j.xplc.2020.100050.

Holmquist M (2000) Alpha/Beta-hydrolase fold enzymes: structures, functions and mechanisms. Curr Protein Pept Sci. 1:209–235. doi: 10.2174/1389203003381405.

Hu J, Liu M, Zhang A, Dai Y, Chen W, Chen F, Wang W, Shen D, Telebanco-Yanoria MJ, Ren B, Zhang H, Zhou H, Zhou B, Wang P, Zhang Z (2022) Co-evolved plant and blast fungus ascorbate oxidases orchestrate the redox state of host apoplast to modulate rice immunity. Mol Plant15:1347–1366. doi: 10.1016/j.molp.2022.07.001.

InterPro; Classification of protein families; European Molecular Biology Laboratory (EMBL) https://www.ebi.ac.uk/interpro/search/sequence/

Iversen R, Andersen PA, Jensen KS, Winther JR, Sigurskjold BW (2010) Thiol-disulfide exchange between glutaredoxin and glutathione. Biochemistry. 49:810–820. doi: 10.1021/bi9015956.

Jaouannet M, Magliano M, Arguel MJ, Gourgues M, Evangelisti E, Abad P, Rosso MN (2013) The root-knot nematode calreticulin Mi-CRT is a key effector in plant defense suppression. Mol Plant Microbe Interact. 26:97–105. doi: 10.1094/MPMI-05-12-0130-R.

Jiménez-Quesada MJ, Traverso JÁ, Alché Jde D (2016) NADPH Oxidase-Dependent Superoxide Production in Plant Reproductive Tissues. Front Plant Sci. 7:359. doi: 10.3389/fpls.2016.00359.

Jing M, Guo B, Li H, Yang B, Wang H, Kong G, Zhao Y, Xu H, Wang Y, Ye W, Dong S, Qiao Y, Tyler BM, Ma W, Wang Y (2016) A Phytophthora sojae effector suppresses endoplasmic reticulum stress-mediated immunity by stabilizing plant Binding immunoglobulin Proteins. Nat Commun. 7:11685. doi: 10.1038/ncomms11685.

Jwa NS, Hwang BK (2017) Convergent Evolution of Pathogen Effectors toward Reactive Oxygen Species Signaling Networks in Plants. Front Plant Sci. 8:1687. doi: 10.3389/fpls.2017.01687.

Kulkarni RD, Kelkar HS, Dean RA (2003) An eight-cysteine-containing CFEM domain unique to a group of fungal membrane proteins. Trends Biochem Sci. 28:118–121. doi: 10.1016/S0968-0004(03)00025-2.

Kunert KJ, Foyer CH (2023). Chapter Three - The ascorbate/glutathione cycle. Advances in Botanical Research 105: 77–112. 10.1016/bs.abr.2022.11.004

Lazar N, Mesarich CH, Petit-Houdenot Y, Talbi N, Li de la Sierra-Gallay I, Zélie E, Blondeau K, Gracy J, Ollivier B, Blaise F, Rouxel T, Balesdent MH, Idnurm A, van Tilbeurgh H, Fudal I (2022) A new family of structurally conserved fungal effectors displays epistatic interactions with plant resistance proteins. PLoS Pathog. 18:e1010664. doi: 10.1371/journal.ppat.1010664.

Le Naour-Vernet M, Lahfa M, Maidment JHR, Padilla A, Roumestand C, de Guillen K, Kroj T, Césari S (2025) Structure-guided insights into the biology of fungal effectors. New Phytol. 246:1460–1477. doi: 10.1111/nph.70075.

Lesage G, Bussey H (2006) Cell wall assembly in Saccharomyces cerevisiae. Microbiol Mol Biol Rev. 70:317–343. doi: 10.1128/MMBR.00038-05.

Li G, Newman M, Yu H, Rashidzade M, Martínez-Soto D, Caicedo A, Allen KS, Ma LJ (2024) Fungal effectors: past, present, and future. Curr Opin Microbiol. 81:102526. doi: 10.1016/j.mib.2024.102526.

Li Y, Wang K, Xie H, Wang YT, Wang DW, Xu CL, Huang X, Wang DS (2015) A Nematode Calreticulin, Rs-CRT, Is a Key Effector in Reproduction and Pathogenicity of Radopholus similis. PLoS One 10:e0129351. doi: 10.1371/journal.pone.0129351.

Lin B, Qing X, Liao J, Zhuo K (2020) Role of Protein Glycosylation in Host-Pathogen Interaction. Cells. 9:1022. doi: 10.3390/cells9041022.

Liu C, Talbot NJ, Chen XL (2021) Protein glycosylation during infection by plant pathogenic fungi. New Phytol. 230:1329–1335. doi: 10.1111/nph.17207.

Liu K, Wang X, Li Y, Shi Y, Ren Y, Wang A, Zhao B, Cheng P, Wang B (2023) Protein Disulfide Isomerase FgEps1 Is a Secreted Virulence Factor in Fusarium graminearum. J Fungi (Basel) 9:1009. doi: 10.3390/jof9101009.

Lo Leggio, L., Simmons, T., Poulsen, JC., et al. (2015) Structure and boosting activity of a starch-degrading lytic polysaccharide monooxygenase. Nat Commun 6: 5961 10.1038/ncomms6961

Lv J, Zhou J, Chang B, Zhang Y, Feng Z, Wei F, Zhao L, Zhang Y, Feng H (2022) Two Metalloproteases VdM35-1 and VdASPF2 from Verticillium dahliae Are Required for Fungal Pathogenicity, Stress Adaptation, and Activating Immune Response of Host. Microbiol Spectr 10:e0247722. doi: 10.1128/spectrum.02477-22

Ma H, Wang M, Gai Y, Fu H, Zhang B, Ruan R, Chung KR, Li H (2018) Thioredoxin and Glutaredoxin Systems Required for Oxidative Stress Resistance, Fungicide Sensitivity, and Virulence of Alternaria alternata. Appl Environ Microbiol. 84:e00086–18. doi: 10.1128/AEM.00086-18.

Malinovsky FG, Fangel JU, Willats WG (2014) The role of the cell wall in plant immunity. Front Plant Sci. 5:178. doi: 10.3389/fpls.2014.00178.

Manzl, C., Enrich, J., Ebner, H., Dallinger, R., Krumschnabel, G. (2004) Copper-induced formation of reactive oxygen species causes cell death and disruption of calcium homeostasis in trout hepatocytes. Toxicology. 196:57–64. doi: 10.1016/j.tox.2003.11.001.

Martínez-Cruz J, Romero D, de la Torre FN, Fernández-Ortuño D, Torés JA, de Vicente A, Pérez-García A (2018) The Functional Characterization of Podosphaera xanthii Candidate Effector Genes Reveals Novel Target Functions for Fungal Pathogenicity. Mol Plant Microbe Interact. 31:914–931. doi: 10.1094/MPMI-12-17-0318-R.

McCabe PM, Van Alfen NK (1999) Secretion of cryparin, a fungal hydrophobin. Appl Environ Microbiol. 65:5431–5435. doi: 10.1128/AEM.65.12.5431-5435.1999.

Mehrabi R, Bahkali AH, Abd-Elsalam KA, Moslem M, Ben M’barek S, Gohari AM, Jashni MK, Stergiopoulos I, Kema GH, de Wit PJ. (2011) Horizontal gene and chromosome transfer in plant pathogenic fungi affecting host range. FEMS Microbiol 35:542–554. doi: 10.1111/j.1574-6976.2010.00263.x.

Mekonen M, Tesfaye K, Mengiste T, Chala A, Nida H, Mekonnen T, Abreha KB, Geleta M (2024) Pathotype determination of Sorghum anthracnose (Colletotrichum sublineola) isolates from Ethiopia using Sorghum differentials. Front Microbiol. 15:1458450. doi: 10.3389/fmicb.2024.1458450.

Münch S, Lingner U, Floss DS, Ludwig N, Sauer N, Deising HB (2008) The hemibiotrophic lifestyle of Colletotrichum species. J Plant Physiol. 165:41–51. doi: 10.1016/j.jplph.2007.06.008.

Münch A, Stingl L, Jung K, Heermann R (2008) Photorhabdus luminescens genes induced upon insect infection. BMC Genomics. 9:229. doi: 10.1186/1471-2164-9-229.

Nakhasi HL, Pogue GP, Duncan RC, Joshi M, Atreya CD, Lee NS, Dwyer DM (1998) Implications of calreticulin function in parasite biology. Parasitol Today 14:157–160. doi: 10.1016/s0169-4758(97)01180-0.

Nielsen H, Tsirigos KD, Brunak S, von Heijne G (2019) A Brief History of Protein Sorting Prediction. Protein J. 38:200–216. doi: 10.1007/s10930-019-09838-3.

Ning N, Xie X, Yu H, Mei J, Li Q, Zuo S, Wu H, Liu W, Li Z (2022) Plant Peroxisome-Targeting Effector MoPtep1 Is Required for the Virulence of Magnaporthe oryzae. Int J Mol Sci 23:2515. doi: 10.3390/ijms23052515.

Njus D, Asmaro K, Li G, Palomino E (2023) Redox cycling of quinones reduced by ascorbic acid. Chem Biol Interact. 373:110397. doi: 10.1016/j.cbi.2023.110397.

Ogura M, Kanesaki Y, Yoshikawa H, Haga K (2025) The DnaJK chaperone of Bacillus subtilis post-transcriptionally regulates gene expression through the YlxR(RnpM)/RNase P complex. mBio. 16:e0405324. doi: 10.1128/mbio.04053-24.

Ohtaki S, Maeda H, Takahashi T, Yamagata Y, Hasegawa F, Gomi K, Nakajima T, Abe K (2006) Novel hydrophobic surface binding protein, HsbA, produced by Aspergillus oryzae. Appl Environ Microbiol. 72:2407–2413. doi: 10.1128/AEM.72.4.2407-2413.2006.

Oide S, Tanaka Y, Watanabe A, Inui M (2019) Carbohydrate-binding property of a cell wall integrity and stress response component (WSC) domain of an alcohol oxidase from the rice blast pathogen Pyricularia oryzae. Enzyme Microb Technol. 125:13–20. doi: 10.1016/j.enzmictec.2019.02.009.

Pan L, Wen S, Yu J, Lu L, Zhu X, Zhang Z (2020) Genome-Wide Identification of M35 Family Metalloproteases in Rhizoctonia cerealis and Functional Analysis of RcMEP2 as a Virulence Factor during the Fungal Infection to Wheat. Int J Mol Sci. 21:2984. doi: 10.3390/ijms21082984.

Parada-Rojas CH, Stahr M, Childs KL, Quesada-Ocampo LM (2024) Effector Repertoire of the Sweetpotato Black Rot Fungal Pathogen Ceratocystis fimbriata. Mol Plant Microbe Interact 37:315–326. doi: 10.1094/MPMI-09-23-0146-FI.

Percudani R, Montanini B, Ottonello S (2005) The anti-HIV cyanovirin-N domain is evolutionarily conserved and occurs as a protein module in eukaryotes. Proteins. 60:670–678. doi: 10.1002/prot.20543.

Pemberton TJ (2006) Identification and comparative analysis of sixteen fungal peptidyl-prolyl cis/trans isomerase repertoires. BMC Genomics 7:244. doi: 10.1186/1471-2164-7-244.

Pierce BC, Agger JW, Zhang Z, Wichmann J, Meyer AS (2017) A comparative study on the activity of fungal lytic polysaccharide monooxygenases for the depolymerization of cellulose in soybean spent flakes. Carbohydr Res. 449:85–94. doi:10.1016/j.carres.2017.07.004.

Punja, Z. K. (2006) Recent developments toward achieving fungal disease resistance in transgenic plants. Canadian Journal of Plant Pathology, 28: S298–S308. 10.1080/07060660609507387

Robin GP, Kleemann J, Neumann U, Cabre L, Dallery JF, Lapalu N, O’Connell RJ (2018) Subcellular Localization Screening of Colletotrichum higginsianum Effector Candidates Identifies Fungal Proteins Targeted to Plant Peroxisomes, Golgi Bodies, and Microtubules. Front Plant Sci. 9:562. doi: 10.3389/fpls.2018.00562.

Rozano L, Jones DAB, Hane JK, Mancera RL (2024) Template-Based Modelling of the Structure of Fungal Effector Proteins. Mol Biotechnol. 66:784–813. doi: 10.1007/s12033-023-00703-4.

Salomon D, Orth K (2013) What pathogens have taught us about posttranslational modifications. Cell Host Microbe.14:269–279. doi: 10.1016/j.chom.2013.07.008.

Samal A, Craig JP, Coradetti ST, Benz JP, Eddy JA, Price ND, Glass NL (2017) Network reconstruction and systems analysis of plant cell wall deconstruction by *Neurospora crassa*. Biotechnol Biofuels. 210:225. doi: 10.1186/s13068-017-0901-2.

Samalova M, Mélida H, Vilaplana F, Bulone V, Soanes DM, Talbot NJ, Gurr SJ (2017) The β-1,3-glucanosyltransferases (Gels) affect the structure of the rice blast fungal cell wall during appressorium-mediated plant infection. Cell Microbiol. 19:e12659. doi: 10.1111/cmi.12659.

Sánchez-Vallet A, Tian H, Rodriguez-Moreno L, Valkenburg DJ, Saleem-Batcha R, Wawra S, Kombrink A, Verhage L, de Jonge R, van Esse HP, Zuccaro A, Croll D, Mesters JR, Thomma BPHJ (2020) A secreted LysM effector protects fungal hyphae through chitin-dependent homodimer polymerization. PLoS Pathog.16:e1008652. doi: 10.1371/journal.ppat.1008652

Scavuzzo-Duggan T, Varoquaux N, Madera M, Vogel JP, Dahlberg J, Hutmacher R, Belcher M, Ortega J, Coleman-Derr D, Lemaux P, Purdom E, Scheller HV (2021) Cell Wall Compositions of Sorghum bicolor Leaves and Roots Remain Relatively Constant Under Drought Conditions. Front Plant Sci.12:747225. doi: 10.3389/fpls.2021.747225.

Seong K and Krasileva KV (2023). Prediction of effector protein structures from fungal phytopathogens enables evolutionary analyses. Nat Microbiol. 8:174–187. doi: 10.1038/s41564-022-01287-6.

Shang S, Liu G, Zhang S, Liang X, Zhang R, Sun G (2024) A fungal CFEM-containing effector targets NPR1 regulator NIMIN2 to suppress plant immunity. Plant Biotechnol J. 22:82–97. doi: 10.1111/pbi.14166.

Shiojima Y, Sano R, Kozono T, Nishikawa A, Kojima Y, Yoshida M, Sunagawa N, Igarashi K, Tonozuka T (2025) Crystal Structure of CcGH131B, a Protein Belonging to Glycoside Hydrolase Family 131 from the Basidiomycete Coprinopsis cinerea. Journal of Applied Glycoscience 72: 7202104. 10.5458/jag.7202104

Singh K, Zouhar M, Mazakova J, Rysanek P (2014) Genome wide identification of the immunophilin gene family in Leptosphaeria maculans: a causal agent of Blackleg disease in Oilseed Rape (Brassica napus). OMICS.18:645–657. doi: 10.1089/omi.2014.0081.

Singh R, Dangol S, Chen Y, Choi J, Cho YS, Lee JE, Choi MO, Jwa NS (2016) *Magnaporthe oryzae* Effector AVR-Pii Helps to Establish Compatibility by Inhibition of the Rice NADP-Malic Enzyme Resulting in Disruption of Oxidative Burst and Host Innate Immunity. Mol Cells. 39:426–438. doi: 10.14348/molcells.2016.0094.

Singh Y, Nair AM, Verma PK (2021) Surviving the odds: From perception to survival of fungal phytopathogens under host-generated oxidative burst. Plant Commun. 2:100142. doi: 10.1016/j.xplc.2021.100142.

Sonah H, Deshmukh RK, Bélanger RR (2016) Computational Prediction of Effector Proteins in Fungi: Opportunities and Challenges. Front Plant Sci.7:126. doi: 10.3389/fpls.2016.00126.

Sperschneider J, Dodds PN (2022) EffectorP 3.0: Prediction of Apoplastic and Cytoplasmic Effectors in Fungi and Oomycetes. Mol Plant Microbe Interact.35:146–156. doi: 10.1094/MPMI-08-21-0201-R.

Sperschneider J, Catanzariti AM, DeBoer K, Petre B, Gardiner DM, Singh KB, Dodds PN, Taylor JM (2017). LOCALIZER: subcellular localization prediction of both plant and effector proteins in the plant cell. Sci Rep. 7:44598. doi: 10.1038/srep44598.

Stolf BS, Smyrnias I, Lopes LR, Vendramin A, Goto H, Laurindo FR, Shah AM, Santos CX (2011) Protein disulfide isomerase and host-pathogen interaction. The Scientific World Journal.11:1749–1761. doi: 10.1100/2011/289182.

Stone LBL, Padilla-Guerrero IE, Bidochka MJ (2022) Fungal Effector Proteins: Molecular Mediators of Fungal Symbionts of Plants. Microbial Cross-talk in the Rhizosphere, pp 297–321; Horwitz BA, Mukherjee PK (eds.) Springer Nature. 10.1007/978-981-16-9507-0_12

Tahir J, Rashid M, Afzal AJ (2019) Post-translational modifications in effectors and plant proteins involved in host–pathogen conflicts. Plant Pathology 68: 628–644. 10.1111/ppa.12983

Tai E-S, Hsieh, P-C, Sheu S-C (2013) Purification and characterization of polygalacturonase from screened Aspergillus tubingensis for coffee processing. Food Sci. Technol. Res., 19:813–818. 10.3136/fstr.19.813

Tamayo E, Benabdellah K, Ferrol N (2016) Characterization of Three New Glutaredoxin Genes in the Arbuscular Mycorrhizal Fungus Rhizophagus irregularis: Putative Role of RiGRX4 and RiGRX5 in Iron Homeostasis. PLoS One.11:e0149606. doi: 10.1371/journal.pone.0149606.

Todd JNA, Carreón-Anguiano KG, Islas-Flores I, Canto-Canché B (2022) Fungal Effectoromics: A World in Constant Evolution. Int J Mol Sci. 23:13433. doi: 10.3390/ijms232113433.

Vasistha P, Singh PP, Srivastava D, Johny L, Shukla S (2025) Effector proteins of Funneliformis mosseae BR221: unravelling plant-fungal interactions through reference-based transcriptome analysis, in vitro validation, and protein‒protein docking studies. BMC Genomics. 26:42. doi: 10.1186/s12864-024-10918-7.

Vela S, Wolf ESA, Rollins JA, Cuevas HE, Vermerris W (2024) Dual-RNA-sequencing to elucidate the interactions between Sorghum and Colletotrichum sublineola. Front Fungal Biol. 5:1437344. doi: 10.3389/ffunb.2024.1437344.

Verna J, Lodder A, Lee K, Vagts A, Ballester R (1997) A family of genes required for maintenance of cell wall integrity and for the stress response in Saccharomyces cerevisiae. Proc Natl Acad Sci U S A. 94:13804–13809. doi: 10.1073/pnas.94.25.13804.

Viaud MC, Balhadère PV, Talbot NJ (2002) A Magnaporthe grisea cyclophilin acts as a virulence determinant during plant infection. Plant Cell.14:917–930. doi: 10.1105/tpc.010389.

Vleeshouwers VGAA, Oliver RP (2014) Effectors as Tools in Disease Resistance Breeding Against Biotrophic, Hemibiotrophic, and Necrotrophic Plant Pathogens. Molecular Plant-Microbe Interactions 27: 196–206. 10.1094/MPMI-10-13-0313-IA

Wan J, He M, Hou Q, Zou L, Yang Y, Wei Y, Chen X (2021) Cell wall associated immunity in plants. Stress Biol. 1:3. doi: 10.1007/s44154-021-00003-4.

Wang Y, Yang Z, Xue J, Wang Y, Li H, Wu Z, Gao Y (2025) Cepharanthine Inhibits Fusarium solani via Oxidative Stress and CFEM Domain-Containing Protein Targeting. Microorganisms.13:1423. doi: 10.3390/microorganisms13061423.

Wang Z, Lienemann M, Qiau M, Linder MB (2010) Mechanisms of protein adhesion on surface films of hydrophobin. Langmuir. 2010 Jun 1;26(11):8491–6. doi: 10.1021/la101240e.

Wawra S, Fesel P, Widmer H, Neumann U, Lahrmann U, Becker S, Hehemann JH, Langen G, Zuccaro A (2019) FGB1 and WSC3 are in planta-induced β-glucan-binding fungal lectins with different functions. New Phytol. 222:1493–1506. doi: 10.1111/nph.15711.

Wharton PS, Julian AM, O’Connell RJ (2001) Ultrastructure of the Infection of Sorghum bicolor by Colletotrichum sublineolum. Phytopathology. 91:149–158. doi: 10.1094/PHYTO.2001.91.2.149.

Wright HT, Sandrasegaram G, Wright CS (1991) Evolution of a family of N-acetylglucosamine binding proteins containing the disulfide-rich domain of wheat germ agglutinin. J Mol Evol. 33: 283–294. 10.1007/BF02100680

Xiao F, Xu W, Hong N, Wang L, Zhang Y, Wang G (2022) A Secreted Lignin Peroxidase Required for Fungal Growth and Virulence and Related to Plant Immune Response. Int J Mol Sci. 23:6066. doi: 10.3390/ijms23116066.

Xu L, Gong W, Zhang H, Perrett S, Jones GW (2018) The same but different: the role of Hsp70 in heat shock response and prion propagation. Prion.12:170–174. doi: 10.1080/19336896.2018.1507579.

Xu Q, Tang C, Wang X, Sun S, Zhao J, Kang Z, Wang X (2019) An effector protein of the wheat stripe rust fungus targets chloroplasts and suppresses chloroplast function. Nat Commun. 10:5571. doi: 10.1038/s41467-019-13487-6.

Yu W, Kang X, Cui X, Hu J, Pan Y, Deng Y, Zhang S (2024) Protein disulfide isomerase MoPdi1 regulates fungal development, virulence, and endoplasmic reticulum homeostasis in Magnaporthe oryzae Journal of Integrative Agriculture, 24: 4670–4689. 10.1016/j.jia.2024.03.054

Zavaliev R and Dong X (2024) NPR1, a key immune regulator for plant survival under biotic and abiotic stresses. Mol Cell. 84:131–141. doi: 10.1016/j.molcel.2023.11.018.

Zeiner M, Niyaz Y, Gehring U (1999) The hsp70-associating protein Hap46 binds to DNA and stimulates transcription. Proc Natl Acad Sci U S A. 96:10194–10199. doi: 10.1073/pnas.96.18.10194.

Zeng T, Rodriguez-Moreno L, Mansurkhodzaev A, Wang P, van den Berg W, Gasciolli V, Cottaz S, Fort S, Thomma BPHJ, Bono JJ, Bisseling T, Limpens E (2020) A lysin motif effector subverts chitin-triggered immunity to facilitate arbuscular mycorrhizal symbiosis. New Phytol. 225:448–460. doi: 10.1111/nph.16245.

Zhang ZN, Wu QY, Zhang GZ, Zhu YY, Murphy RW, Liu Z, Zou CG (2015) Systematic analyses reveal uniqueness and origin of the CFEM domain in fungi. Sci Rep. 5:13032. doi: 10.1038/srep13032.

